# A balance between matrix deformation and the coordination of turning events governs directed neutrophil migration in 3-D matrices

**DOI:** 10.1101/2020.09.02.868505

**Authors:** Joshua François, Adithan Kandasamy, Yi-Ting Yeh, Cindy Ayala, Ruedi Meili, Shu Chien, Juan C. Lasheras, Juan C. del Álamo

**Author notes:** corresponding authors: Juan C. del Álamo, Juan C. Lasheras, Joshua François.

## Abstract

Three-dimensional (3-D) neutrophil migration is essential for immune surveillance and inflammatory responses. During 3-D migration, especially through extravascular spaces, neutrophils rely on frontal protrusions and rear contractions to squeeze and maneuver through extracellular matrices containing narrow pores. However, the role of matrix density and the cells’ ability to probe and remodel matrix pores during 3-D chemotaxis are far from being understood. We investigated these processes by tracking the trajectories of over 20,000 neutrophils in a 3-D migration device containing collagen matrices of varying concentrations and analyzing the shape of these trajectories at multiple scales. Additionally, we quantified the transient 3-D matrix deformations caused by the migrating cells. The mean pore size of our reconstituted collagen matrices decreased when the collagen concentration ([col]) was increased. In low-[col] matrices, neutrophils exerted large transient deformations and migrated in relatively straight trajectories. In contrast, they were not able to appreciably deform high- [col] matrices and adapted to this inability by turning more often to circumvent these narrow matrix pores. While this adaptation resulted in slower migration, the cells were able to balance the more frequent turning with the long-range directional bias necessary for chemotaxis. Based on our statistical analysis of cell trajectories, we postulate that neutrophils achieve this balance by using matrix obstacles as pivoting points to steer their motion towards the chemoattractant. Inhibiting myosin-II contractility or Arp2/3-mediated pseudopod protrusions not only compromised the cells’ ability to deform the matrix, but also made them switch to increased turning in more restrictive matrices when compared to untreated control cells. Both myosin-II contractility and Arp2/3-mediated branched polymerization of actin played a role in fast migration, but Arp2/3 was also crucial for neutrophils when coordinating the orientations of successive turns to prevent veering away from the chemotactic path. These results may contribute to an improved understanding of the mechanisms employed by migrating neutrophils in confined 3-D environments, as well as the molecular and environmental regulators for maintaining persistent motion.

## 1. Introduction

Three-dimensional (3-D) cell migration is an important process during embryonic development, angiogenesis, cancer cell metastasis, immune surveillance, and inflammatory responses [1-5]. During inflammatory responses, neutrophils, which are a subtype of white blood cells (leukocytes) and the first responders of the innate immune system, travel along chemotactic gradients initiated at or near sites of injury or infection [6]. This journey involves neutrophil activation, rolling and migration within post-capillary blood vessels, transmigration across vessel walls, and 3-D migration through the extravascular space guided by chemokine gradients [5]. During these processes, neutrophils must negotiate a wide distribution of pore sizes and densities in order to translocate [7]. Migration in the 3-D extravascular space is particularly complex, as it requires navigating through crowded spaces composed of other cells, glycoproteins, and extracellular matrix (ECM) proteins [8, 9].

To explain how neutrophils migrate through 3-D extravascular spaces, previous studies have pointed to two main maneuvers: (1) frontal protrusions and (2) rear contractions [8, 10]. Frontal protrusions are typically in the form of lamellipodia, which are an arrangement of rosettes formed by multiple, interleaved thin lamella [11]. These structures form as a result of branched polymerization of actin at the front of polarized cells, which is in turn mediated by a set of actin binding proteins known as the Arp2/3 complex [12, 13]. The Arp2/3 complex plays an important role in path finding during 3-D leukocyte migration [11]. Specifically, the protein complex facilitates lamellipodial formation for environmental probing and helps cells find neighboring pores large enough to squeeze through. Rear contractions are mediated by non-muscle myosin-II, a motor protein that is important for cytokinesis, organelle transport, and cell migration [14, 15]. Non-muscle myosin-II has long been appreciated for its role in cell migration, and has recently been found to play an important role in 3-D leukocyte migration in restrictive microenvironments [16].

Efficient path-finding depends on the ability of motile cells to change directions before committing to a path [17]. After deciding on a path, cells must then translocate their bodies, including their bulky nucleus, through pores while still maintaining their compass aligned with the chemotactic gradient [18]. To perform this task, neutrophils combine traction forces, which help them squeeze through and enlarge matrix pores, with turning to circumvent impassable pores [18, 19]. These two mechanisms are challenged by increasing collagen concentration, which would decrease both pore size and matrix deformability. Yet, neutrophils are able to exhibit directed migration in a wide variety of extracellular environments, including the highly restrictive ones [20].

Separately, previous studies have examined the effect of collagen density on 3-D cell migration, providing insight into the relationship between changes in cell kinematics and surrounding extracellular matrices [16, 21-25]. However, many of these studies only considered randomly migrating cells and analyzed their motion at different timescales; this makes it difficult to separate cell speed from trajectory shape. Moreover, many of these studies focused on cancer cells, which are able to secrete matrix metalloproteinases to degrade the extracellular matrix material. Matrix degradation reduces the need for cells to bypass impenetrable sections along a migratory path (e.g. narrow pores), and therefore enables relatively straight trajectories [24]. Findings from these cancer-cell-based studies may therefore not be fully applicable to contexts where degradation is reduced or not possible. Finally, most previous works did not quantify 3-D matrix deformations induced by traction forces. The present manuscript addresses these gaps in knowledge by analyzing the trajectories of chemotactic neutrophils at different length scales, with a special emphasis on how these cells combine physical forces and turning events to maintain directional motion towards the chemoattractant source.

## 2. Materials and Methods

### 2.1 Cell culture, differentiation, and cytoplasmic labeling

We cultured and differentiated cells from the human promyelocytic leukemia cell line (HL-60, ATCC) into neutrophil-like cells (dHL-60) as previously described [26]. We grew HL-60 cells to approximately 1 × 10^6^ cells/ml and passaged the cells every 2-3 days in Roswell Park Memorial Institute medium (RPMI-1640, Thermo Fisher Scientific) supplemented with L-glutamine and 10% fetal bovine serum (FBS, Omega Scientific). We differentiated HL-60 cells by taking 1.5 × 10^6^ cells from each passage and placing them in supplemented RPMI-1640 medium with the additional 1.3% dimethyl sulfoxide (DMSO, Sigma). Next, we incubated Both HL-60 and dHL-60 cells at 37° C and 5% CO_2_. We then performed experiments with dHL-60 cells 4 days after being cultured in the presence of DMSO. For experiments involving fluorescently labeled cells, we labeled day 4 differentiated dHL-60 cells with CellTracker Deep Red (Thermo Fisher Scientific) prior to mixing them with collagen solutions. Cells were incubated in a 10 nM solution of CellTracker in RPMI-1640 without FBS for 30 minutes at 37° C and 5% CO_2_. Afterwards, the cells were centrifuged and re-suspended in RPMI-1640 containing 10% FBS.

### 2.2 *In-vitro* 3-D directed migration assay

We adapted a version of a previously published custom-built chemotaxis device to study 3-D neutrophil chemotaxis (Figure 1a) [27, 28]. First, we treated 25-mm glass coverslips (Fisher Scientific) on one side with 250 μl of 0.1-M NaOH for 5 minutes. We then removed the NaOH and rinsed the coverslips with distilled H_2_O. We let the coverslips dry and added (3-aminopropyl) triethoxysilane (APTES, Sigma) to their treated sides afterwards. After 30 minutes, we rinsed the coverslips again with distilled H_2_O and placed them on Kimwipes™ for air-drying, with the treated surfaces face up. Next, we punched 12-mm diameter holes in the center of 50×9-mm Falcon petri dishes (BD Falcon). We then attached treated glass coverslips to the bottoms of the petri dishes with their treated sides face up using vacuum grease (Beckman Vacuum Grease Silicone). We attached separate treated glass coverslips to the tops of the holes with their treated sides face down and covering the majority of the holes’ diameters. This sandwiching of coverslips created a pocket that was used later to place collagen gel solutions. Before fabricating collagen gels, we rested the devices on top of a metal block that sat in a container filled with ice.

**Figure 1.**
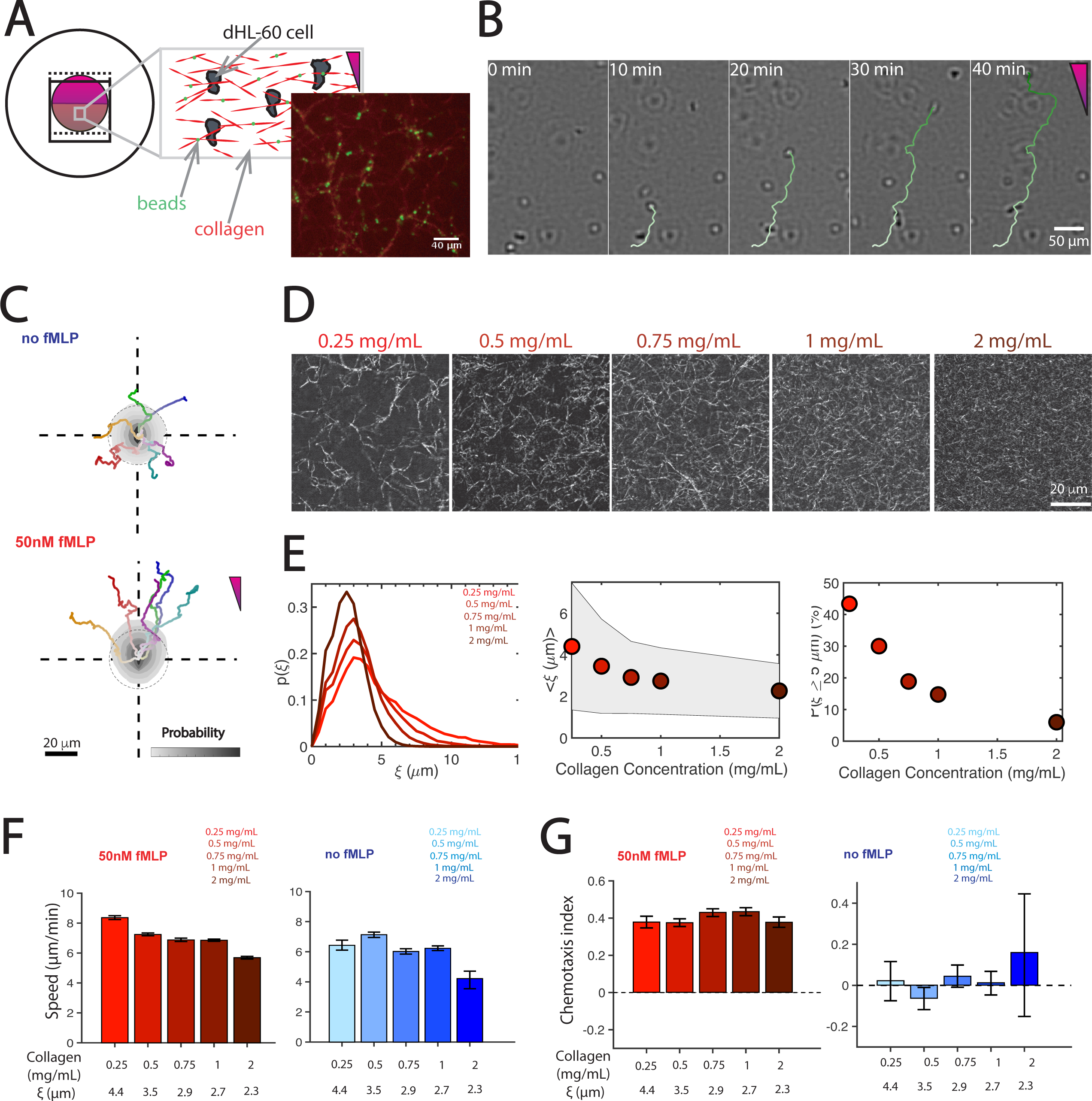
Experimental Setup and Main Migratory Statistics. **A)** Schematic of custom 3-D migration chamber with dHL-60 cells (gray) and fluorescent micro-beads (green) embedded inside of a rat-tail type I collagen gel (red). Imaging of fluorescently labeled collagen fibers showed that micro-beads are localized to collagen fibers after gel formation. **B)** Brightfield image sequence of a cell chemotaxing along an fMLP gradient, and automated tracking of the cell trajectory (green). **C)** Probability density maps of cell coordinates (gray contours), and representative trajectories of dHL-60 cells in the absence (top) and presence (bottom) of fMLP gradient. The data come from 0.75 mg/ml gels. The dashed circles represent unbiased random motion. In panels **B** and **C**, time progression along each trajectory is represented by changes from white to saturated colors, and the direction of the fMLP gradient is indicated by purple triangle. **D)** Confocal reflection microscopy images of extracellular matrices with collagen concentrations ranging from 0.25 - 2 mg/ml. **E)** <u>Left panel:</u> Probability distributions of matrix pore size radius ξ from bubble analysis of reflection images. <u>Center panel:</u> Mean pore size ⟨ξ⟩ vs. collagen concentration. <u>Right panel:</u> Fraction of available pores with ⟨ξ⟩ ≥5 μm vs. collagen concentration. **F)** Mean cell speeds vs. matrix density (expressed as collagen concentration or ⟨ξ⟩) in the presence (left) and absence (right) of fMLP gradient. **G)** Chemotactic index vs. matrix density in the presence (left) and absence (right) of fMLP gradient. Error bars represent 95% confidence interval.

### 2.3 Collagen gel fabrication and fluorescent labeling

We fabricated collagen gels by neutralizing solutions of rat-tail type-I collagen dissolved in 0.2-N acetic acid (Corning) with 1-N NaOH in proportions recommended by the manufacturer. We added a suspension of supplemented RPMI 1640 medium containing 4.7×10^3^ dHL-60 cells instead of the recommended 10X PBS and water. We also mixed a 1.2% solution of fluorescent microspheres (Molecular Probes) with the collagen solution. While mixing the different constituents of the collagen gel, we placed all solutions in a small container filled with ice. After the collagen gel solution was made, we pipetted 113 μl of the solution into the pocket of each device and placed each device in an incubator at 37° C and 5% CO_2_ for 30 minutes. We placed the devices in an incubator vertically to allow the gel solution to settle to the lower half of the pocket. For experiments involving reflection microscopy imaging, we fabricated collagen gels in the same manner as described above, leaving out the cells and fluorescent microspheres in the supplemented RPMI 1640 medium. We placed 100 μl of the gel solution on top of glass coverslips attached to the bottoms of petri dishes with 12-mm diameter holes. We did not add a second coverslip on top of the holes. We then put each petri dish in an incubator at 37° C and 5% CO_2_ for 30 minutes. Afterwards, we filled these dishes with 2 ml of supplemented RPMI 1640.

We fluorescently labeled collagen fibers with 5-Carboxytetramethylrhodamine (TAMRA, ThermoFisher) by following a previously published protocol [29]. We first added 25 mg of TAMRA to a 2.5 ml solution of DMSO and stored at −20° C in an Eppendorf tube. We covered the Eppendorf tube with aluminum foil to protect the solution from light. Next, we made a labeling buffer of 0.25 M NaHCO_3_ and 0.4 M NaCl, and subsequently adjusted the pH to 9.5 by using a 10 M solution of NaOH. We then filled a 1 ml syringe with an 8.46 mg/ml solution of rat-tail type I collagen in a 0.2 N acetic acid solution (Corning) and injected the solution into a 3 ml dialysis cassette. The cassette we used had a 10 kDa molecular weight cutoff. Before retracting the hypodermic needle, we removed air from the cassette by pulling the syringe plunger up. We then performed an overnight dialysis in a beaker containing a 1 l solution of our labeling buffer at 4° C. 100 μl of our 10 mg/ml TAMRA solution was mixed with 900 μl of our labeling buffer, with both solutions at room temperature. After mixing, we placed the TAMRA solution in a 4° C fridge. The collagen was removed from the cassette using a 2 ml syringe and 1 ml of the solution was mixed with 1 ml of diluted TAMRA using a pipette. We then placed the resulting solution in a micro-centrifuge tube and incubated with rotation overnight at 4° C. At this point, the collagen was labeled with the TAMRA dye and we put this mix in a dialysis cassette and dialyzed it overnight at 4° C against a 1 l solution of our labeling buffer. Finally, we dialyzed the TAMRA-labeled collagen overnight at 4° C against a 0.2% acetic acid solution with a pH of 4. We used the final volume to calculate the concentration of the TAMRA-labeled collagen.

### 2.4 Inhibition drug treatment experiments

We performed Arp2/3 complex and myosin-II inhibition experiments using ck666 (Sigma) and blebbistatin (Abcam). For each treatment condition, we incubated day 4 differentiated cells with either a 100-μM solution of ck666 in supplanted RMPI, a 10-μM solution of blebbistatin in supplemented RPMI, or both in supplemented RPMI at 37° C and 5% CO_2_ for 30 minutes. We then centrifuged the treated cells and re-suspended the cells in supplemented RPMI medium. Afterwards, we followed the aforementioned protocol for embedding cells in collagen gels, with the addition of ck666, blebbistatin or both so that the final concentration of each gel in these experiments contained 100-μM of ck666 and 10-μM of blebbistatin. We added supplanted medium to the chemotaxis chambers after gel formation. Our chemoattractant solutions containing fMLP also contained the drugs at the same concentrations as those used when preparing the collagen gels.

### 2.5 Imaging

We imaged collagen fibers for pore size analysis using a Leica SP5 microscope in reflection mode with a 40x immersion lens at 2x zoom. We obtained bright-field images for population migration experiments using an enclosed Leica DMI6000 B microscope with a 5x air lens at 37° C and 5% CO_2_. We retracted the microscope’s magnifier to produce images that made automated cell tracking easier. In each experiment, we acquired four planes at 200-μm spacing in each collagen gel every Δ*t* = 30 seconds for 1 hour. We imaged four planes in order to follow a larger a number of cells and we chose 200-μm spacing to avoid imaging the same cells in more than one plane.

We imaged single cells and matrix deformations on an enclosed Zeiss LSM 880 Laser Scanning Microscope in the fast Airyscan mode, with a 40x water lens and 1.3x zoom. We acquired two 80-μm imaging stacks with 0.7-μm spacing between planes every 30 seconds. The duration of time-lapse experiments varied. We imaged cells in each experiment until cells of interest left the field of view. Given that dHL-60 cells are relatively fast-moving cells, we switched imaging channels for the labeled cells and microspheres per line scan during acquisition. This was to minimize significant differences in cell positions with respect to matrix deformations during the acquisition of the two stacks for a single time point. The raw Carl Zeiss Image (CZI) Airyscan output files from the Zeiss LSM 880 microscope underwent Airyscan processing using Zeiss’s imaging software Zen Black. After Airyscan preprocessing, these files were loaded into FIJI [30] and then exported as uncompressed .TIFF files. The .TIFF files were loaded into MATLAB [31] for further quantitative analysis.

### 2.6 Pore size analysis

We used a bubble analysis method to characterize the distribution of pore sizes from confocal reflection images of collagen networks, as previously described [32, 33]. Before performing this analysis, we preprocessed the images in FIJI. First, we removed a radial intensity gradient that appeared as an artifact from our imaging system using FIJI’s sliding paraboloid feature with a radius of 50 pixels under the software’s background subtraction option. We then removed isolated pixels in the images using FIJI’s de-speckle feature, which applied a median filter to the images. Finally, we exported the preprocessed images to MATLAB to perform the bubble analysis.

In our bubble analysis, we binarized each image by setting an intensity threshold equal to the mean image intensity, which resulted in the segmentation of the collagen fibers. We then removed segmented objects containing less than 400 pixels to eliminate noise. Next, we computed the distance to the nearest segmented fiber for all points inside the pores. Afterwards, the resulting distance maps were smoothed out with a Gaussian window of size 100×100 pixels and 1/e-radius of 50 pixels. We identified pore centers by using the local maxima in the distance map, with maximum distance values representing pore sizes. We characterized the pore size distributions in terms of their probability density function *p*(*ξ*), mean pore size ⟨*ξ*⟩, standard deviation, and the probability of finding pores with diameters larger than 5 microns, i.e., *p*(*ξ*≥ 5 *μm*).

### 2.7 Automated cell tracking

We implemented a custom MATLAB algorithm to automatically track cells in time-lapse sequences of bright-field images, *B*(*x,y,t*) (Figure 1B). First, we processed the initial image of each sequence, *B*(*x,y,0*), in 2 steps to identify the positions of in-focus cells.

#### Step 1. Noise removal and segmentation

Since retracting the microscope magnifier produced images where in-focus cells resembled a negative Laplacian of a Gaussian, i.e.,

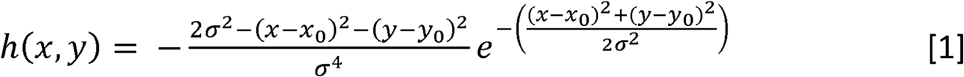

where (*x*_*0*_, *y*_*0*_) are the coordinates of the cell center and *σ* is a free parameter related to cell size, we convolved each image with a centered 6×6-pixel *h*(*x,y*) window. This operation, i.e., 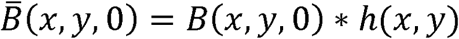 resulted in noise removal. Next, we computed a histogram of image intensities and used intensities higher than the distribution’s first peak, which corresponded to background, as a global threshold to segment cells in 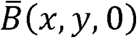.

#### Step 2. Calculation of Cell Positions

We centered a 10×10-pixel search window at the centroid of each cell segmented in Step 1, cropped, and normalized the image by its maximum intensity. We then fitted *h*(*x,y*) to the resulting image using least squares with *x*_*0*_, *y*_*0*_ and *σ* as fit parameters. This operation yielded the coordinates of the center of each cell, 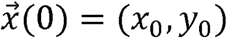.

For subsequent time points, we followed the same 2-step procedure described above with a slight modification that allowed us to track cell trajectories: in step 2, the search window for each cell in frame *t* was centered at the nearest cell centroid determined in the previous frame, i.e., 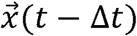.

#### Spurious track removal

We encountered two situations leading to spurious cell tracks. Cells that moved outside of the field of view were lost causing the algorithm to track random points. This resulted in object displacements much larger than the average cell displacements, which were < 20 pixels. Thus, objects that had displacements greater than this threshold between any two successive frames were automatically excluded. Cells that moved too close to one another sometime led algorithm to switch their track labels. Given that these events were not frequent and that, in some cases, it was not straightforward to determine the correct labeling, all cell track pairs that intersected each other at the same time instant were automatically excluded.

#### Stage drift correction

We noted that each experiment contained either dead cells or cells that were non-motile and tracked the motion of these cells as a surrogate of stage drift. Thus, we computed the net displacements of all tracked cells, 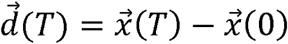, where *T = 60* minutes. We then selected the cells in the bottom 15% of 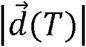 as non-motile cells. Their trajectories were fitted to the model 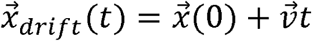, where 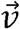 represents the drift velocity, and ensemble averaged over all the non-motile cells, 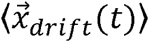. Since drift was systematically largest within the first 20 minutes after adding buffer or fMLP, we did not include data from that initial 20-minute period in our analysis. Drift was removed with 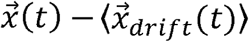, where 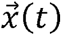 was the trajectory for a cell in the upper 85% of 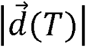 the bracket operator denotes ensemble averaging for all the motile cells.

### 2.8 Analysis of cell trajectories in intrinsic coordinates

We analyzed the geometry of each cell’s trajectory in intrinsic coordinates (i.e., distance traveled, and angular orientation). To perform this analysis, we first tracked cell positions vs. time as described above. We then computed the distance traveled by the cell vs. time, *i*.*e*., 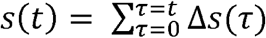, where 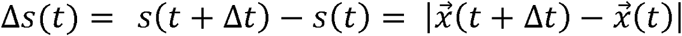, and used this information to obtain 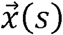 via interpolation. The vector tangent to the trajectory was obtained as 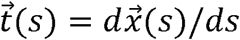. The angular orientation of cell motion with respect to the chemoattractant gradient was quantified by the angle 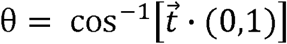. The average cell speed was defined from the length *s* of the cell trajectory over *T = 60* minutes, *i*.*e*., 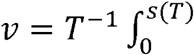. A cell was considered to be motile if its net displacement 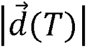 was larger than 30 μm. We computed the chemotaxis index of the trajectories using 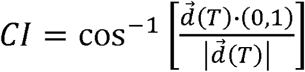. The mean squared displacements (MSD) of the cells were computed as a function of distance lags, *Δs*, i.e., 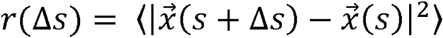, where the bracket operator denotes ensemble averaging for all the motile cells in each experiment and along all values of *s* along each trajectory. Similarly, we computed the autocorrelation function of the vector tangent to the trajectory, 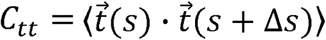. We estimated the persistence distance *S*_*p*_ of the trajectories for each experimental condition by assuming that this correlation decays exponentially with distance traveled, i.e., 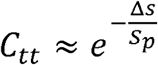. Specifically, we determined the value Δ*s*_0_ for which the correlation function reached the cutoff of *c*_*tt*,0_ = 0.05, and estimated the persistence length as *S*_*p*_ ≈ −Δ_0_/*log C*_*tt*,0_. To better understand the changes in direction along the cells’ trajectories, we calculated the changes in orientation of 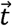 versus *Δs*, i.e., 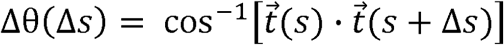, and determined their probability density function (*p*.*d*.*f*.) conditioned to *Δs*, p(θ | Δ*s*). We also determined the joint *p*.*d*.*f*. of speed and changes in orientation, p(θ, *v* | Δ*s*), as well as the p.d.f. of Δθ(Δ*s*) conditioned to θ and Δ*s*, p(Δθ |θ,Δ*s*).

### 2.9. Three-dimensional matrix incremental deformation fields and contractile moments

We computed the incremental changes in 3-D deformation fields inside our collagen gels, 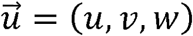, by using a multigrid Particle Image Velocity (PIV) approach [34] that was an amended version of a previous in-house 3-D PIV algorithm [35]. Briefly, we performed a first PIV pass on each confocal stack of fluorescently labeled beads using the stack from the immediately preceding time point as a reference. An interrogation volume of 56×56×24 pixels with a spacing of 28, 28, and 22 pixels respectively, was chosen during this pass. We then ran PIV again using interrogation volumes of 28×28×22 pixels with a spacing of 14, 14, and 11 pixels, respectively. During the second pass, the smaller interrogation windows of the reference stack were shifted by the displacements calculated in the first pass. This resulted in a 3-D deformation field that captured large deformations with the relatively large interrogation volumes in the first pass, while also achieving higher spatial resolution from the smaller interrogation volumes used during the second pass [36]. We constructed visualizations of the 3-D cell shapes and incremental deformation fields by creating Visualization Toolkit (VTK) files, which we imported into the open source software Paraview [37] for 3-D visualization.

To quantify a cell’s ability to deform the matrix, we expanded Butler et al.’s concept of contractile moment to three dimensions and determined the first-order moments of 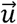 [38]. Using the 3-D cell shape data from confocal stacks of fluorescently labeled cells, we calculated the cell centroids 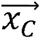 and defined a region around each cell, 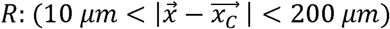, to calculate the moment matrix

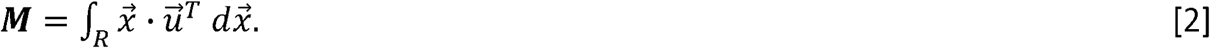

The diagonal elements of *M* provided the total contraction or extension of the collagen gel around individual cells, in each ordinate direction between two consecutive timepoints (Δ*t* = 0.5 min). The net 3-D contractile moment displacing the collagen gel was given by *tr*(***M***) and the time-averaged absolute quantity ⟨ |*tr*(***M***)|⟩ represented a concise metric for gauging cells’ ability to incrementally deform surrounding extracellular matrix fibers.

## 3. Results

### 3.1. Automated analysis of neutrophil chemotaxis in 3-D matrices

We assembled a 3-D migration chamber using a well-established design (See Methods and Figure 1A) [27, 28]. We used type I collagen in our chamber because collagen is the extracellular matrix protein most commonly found in mammals and, specifically, type I collagen is the most common type of collagen [15, 39]. We used neutrophil-like dHL-60 cells differentiated from the human promyelocytic leukemia cell line (HL-60, ATCC). We used fluorescent microbeads attached to the collagen fibers to track matrix deformation (Figure 1A, inset). We chose fMLP – a small peptide secreted by bacteria after infection [40] – as a chemoattractant.

To study the kinematics of directed 3-D neutrophil migration with high throughput, we developed a fully automated algorithm to track cell trajectories from brightfield microscopy time-lapse sequences (Figure 1B). Overall, we tracked 21,150 cell trajectories under different conditions (see Table 1 in the Supplementary Information for details). We confirmed that our system enabled directed neutrophil migration by performing experiments with and without the addition of fMLP. In both conditions, we mapped the probability density *p*(*x,y*) of the cell coordinates in the directions parallel (y) and perpendicular (x) to the chemoattractant gradient, and plotted the 6 longest cell trajectories (Figure 1C). In the absence of fMLP, the distribution of cell trajectories was symmetric leading to circular contours in *p*(*x,y*), as expected for random migration. In contrast, the chemoattractant gradient created by adding fMLP resulted in cell trajectories preferentially directed towards the chemoattractant source, with a *p*(*x,y*) being markedly asymmetric and shifted towards the positive y direction. These data confirm that our system recapitulated 3-D neutrophil chemotaxis, and that our cell-tracking algorithm was able to capture this process from brightfield microscopy, without the need for fluorescent labeling.

### 3.2. Collagen density affects the cell speed but not the chemotaxis of 3-D neutrophils

In physiological contexts, neutrophils must travel across diverse microenvironments. To study the effect of collagen density ([col]) on the ability of neutrophils to undergo directed migration, we fabricated gels with 0.25 mg/ml ≤ [col] ≤ 2.0 mg/ml. Varying collagen concentration has been previously shown to affect the matrix microenvironment, most notably its pore size [21, 41, 42]. Thus, we visualized the structure of our collagen gels (Figure 1D) and quantified their distribution of pore radii (*ξ*) (Figure 1E). This distribution became narrower and shifted to lower values of *ξ* with increasing [col] (Figure 1E, left panel). In particular, ⟨ξ⟩ decreased monotonically from 4.4 to 2.3 *μm* when [col] was increased from 0.25 to 2.0 mg/ml (Figure 1E, center panel). We took *ξ*∼5 *μm* as an approximate value for the radius of dHL-60 cell nuclei [43] and calculated *p*(*ξ* ≥5 *μm*) to assess the availability of pores through which cells could easily squeeze through without significant nuclear deformation. This calculation showed that the fraction of pores larger than *ξ*∼5 *μm* was over 40% in 0.25 mg/ml matrices, but it decreased to under 10% as the [col] increased to 2 mg/ml (Figure 1E, right panel). Overall, these data highlight that the different collagen concentrations considered in our experiments allowed us to study 3-D neutrophil-like migration through microenvironments that ranged from non-restrictive to highly restrictive.

Next, we were interested in how these different matrix microenvironments affected directed 3-D neutrophil migration. We found that cells migrated with mean speeds that decreased from ⟨*v*⟩ = 8.4 μm/min to ⟨*v*⟩ = 5.7 μm/min as [col] was increased and ⟨*ξ*⟩ decreased (Figure 1F, left panel). A similar dependence of ⟨*v*⟩ on ⟨*ξ*⟩ was observed in randomly migrating cells (Figure 1F, right panel). In spite of migrating more slowly as [col] was increased, cells were still able to undergo directed migration towards the fMLP source in all matrices investigated (Figure 1G). Overall, these results indicate that decreasing matrix pore size and availability has a marked effect on the migration speed of wild-type neutrophil-like cells, but not on their ability to perform chemotaxis.

### 3.3. 3-D neutrophil migration requires turning but not mechanical forces to negotiate restrictive matrices

In addition to reducing pore size, increasing [col] from 0.25 to 2 mg/ml is known to cause significant matrix stiffening (i.e., bulk Young’s modulus raised from 440 to 5200 Pa, [42]). Thus, we were interested in finding whether changes in [col] affected the mechanical interactions between neutrophils and their environment. To this end, we imaged fluorescently labeled, chemotaxing cells in our custom-made devices along with fluorescent micro-beads that were attached to matrix fibers. We used Airyscan confocal microscopy to image both the cells and micro-beads in 0.25 and 2 mg/ml collagen gels (Figure 2A, left panel). Matrix deformations were evident in 0.25 mg/ml gels by comparing the location of beads at any given time-point (green) with that at the previous time-point (orange). The 3-D incremental matrix deformations 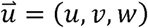 were quantified using PIV (Figure 2A). Our measurements showed clear deformations in the 0.25 mg/ml matrices, with a magnitude (∼3 μm) comparable to the cell nucleus size. These data suggest that, in addition to navigating the matrix searching for large pores, cells in low-[col] matrices are able to significantly deform and enlarge matrix pores (Figure 2B and Supplemental movie 1). In contrast, cells in high-[col] matrices did not exert forces large enough to cause appreciable matrix deformations (Figure 2C and Supplemental movie 2). These observations were confirmed by our statistical analysis of 3-D contraction moments generated by neutrophils for [col] = 0.25, 0.75 and 2 mg/ml (Figure 2D). By extension of Butler et al.’s two-dimensional definition [38], the contraction moments were defined as |*tr* (***M***)|, where the elements of matrix ***M*** are the first-order moments of vector 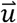 in a sphere of 200 microns surrounding the cell (see eq. 2 in the Methods Section). The statistics of |*tr* (***M***)| indicated a 2.5-fold decrease in matrix deformations between [col] = 0.25 and 0.75 mg/ml, and a 10-fold decrease between [col] = 0.25 and 2 mg/ml.

**Figure 2.**
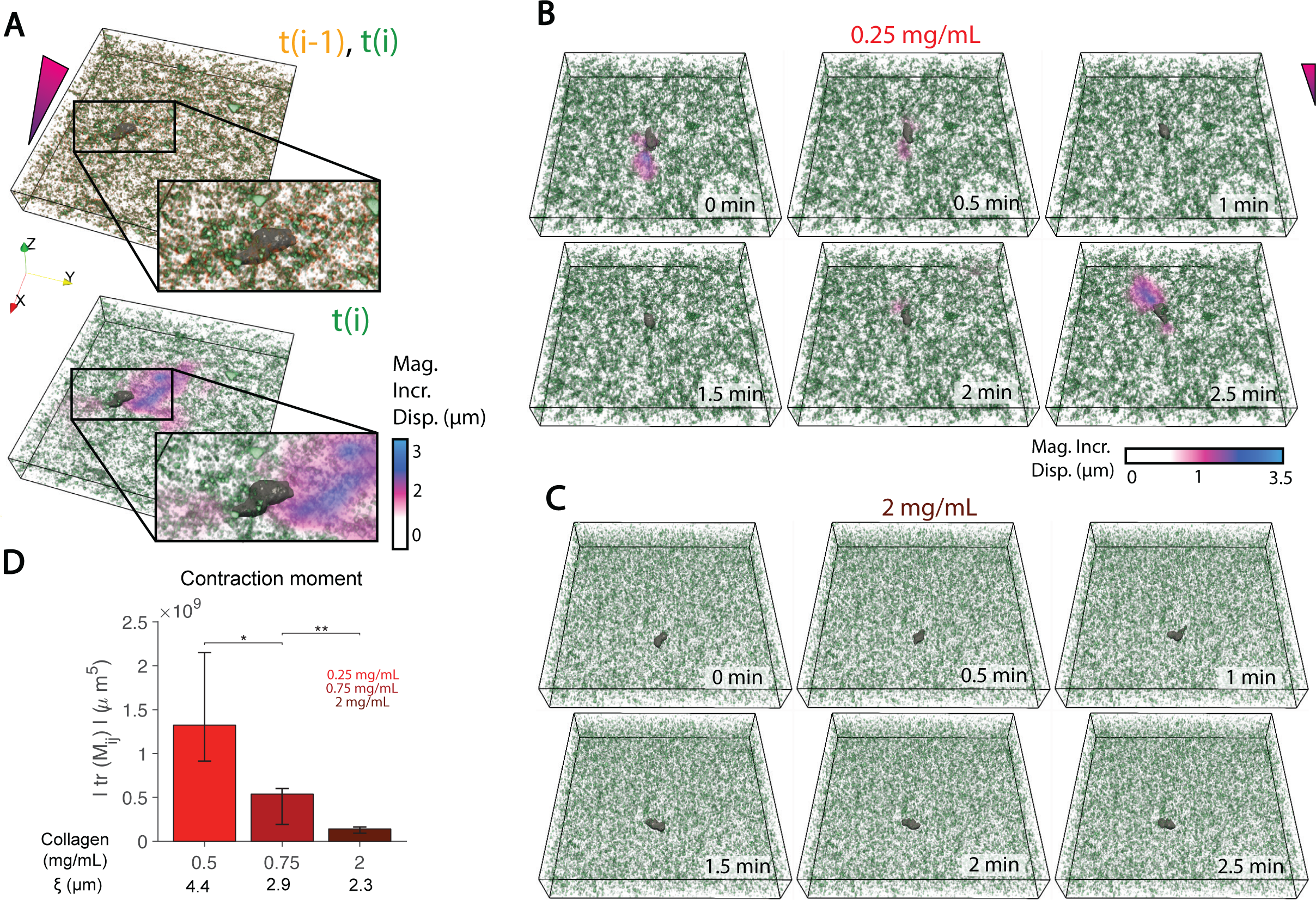
Mechanical Remodeling of the Matrix for Varying Collagen Densities. **A)** <u>Top:</u> Reconstruction of a fluorescently labeled cell (grey), and fluorescent micro-beads at *t* = (*i* − 1)Δ*t* for beads (orange) and *t* = *i*Δ *t* (green) during chemotactic migration in 0.75 mg/ml collagen gel; Δ*t* = 30 seconds. <u>Bottom:</u> Absolute magnitude of incremental deformations computed by 3-D Particle Image Velocimetry. **B)** Representative time-lapse reconstructions of cells (grey), beads (green), and incremental 3-D matrix deformation (magenta – blue colormap) during migration in a low collagen concentration matrix (0.25 mg/ml). **C)** Same as panel **B** for high collagen concentration matrix (2 mg/ml). **D)** Mean magnitude of 3-D contractile moment, |*tr* (***M***)|, vs. collagen density. Error bars represent 95% confidence intervals of the mean. * and ** respectively indicate p<0.05 and p<0.01 according to the Mann-Whitney U test.

Of note, the representative cell in the 2.0 mg/ml collagen matrix initially moved towards the chemoattractant source, but seemed unable to continue moving in that direction at t = 1 min. Subsequently, it performed a sharp, ∼π/2-radian turn and continued to migrate in the direction perpendicular to the fMLP gradient for the next minute. In comparison, the representative cell in the 0.25 mg/ml matrix was able to sustain directed motion towards the fMLP source for t = 2.5 minutes. These observations support the idea that as [col] increases, these cells must adapt their migration strategy to circumvent the more frequent impassable, non-deformable sections of the matrix.

### 3.4. Neutrophils migrating in 3-D retain long-term directional memory despite turning more frequently as matrix pore size decreases

The results presented above (Sections 3.2 and 3.3) led us to hypothesize that turning events are a major factor that enabled the directed migration of neutrophils in restrictive 3-D matrices at the expense of decreasing speed. To examine this hypothesis, we aimed to statistically characterize the cell trajectories in matrices with different collagen concentrations. We first examined the probability distribution *p*(*x,y*) of cell coordinates along their trajectories for different collagen concentrations (Figure 3A). In all the matrices investigated, the trajectories were biased towards the chemoattractant source, and the maps of *p*(*x,y*) had eccentric, elongated contours in the *y* direction. However, the trajectories were shorter and the *p*(*x,y*) contours extended less into the *y*>*0* plane as [col] increased, in agreement with the decreasing cell speed (Figure 1F). Qualitative inspection of the 6 longest trajectories suggested that the cells perform sharper turns with increasing [col] (see arrows in Figure 3A). To characterize turning events statistically, we computed the autocorrelation function of the vector tangent to the trajectory in intrinsic coordinates, 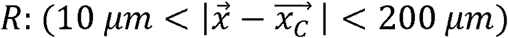 (see diagram in Figure 3B). This function was plotted in Figure 3C for chemotaxing and randomly moving cells in different matrices, showing that *C*_*tt*_ decayed with increasing Δ*s* from its value of *C*_*tt*_ (0) = 1 at the origin as the cells added turns to their trajectories. For chemotaxing cells, this decrease was more pronounced in high-[col] matrices than in low-[col] ones. By contrast, randomly moving cells showed a weak dependence of *C*_*tt*_ with [col].

**Figure 3.**
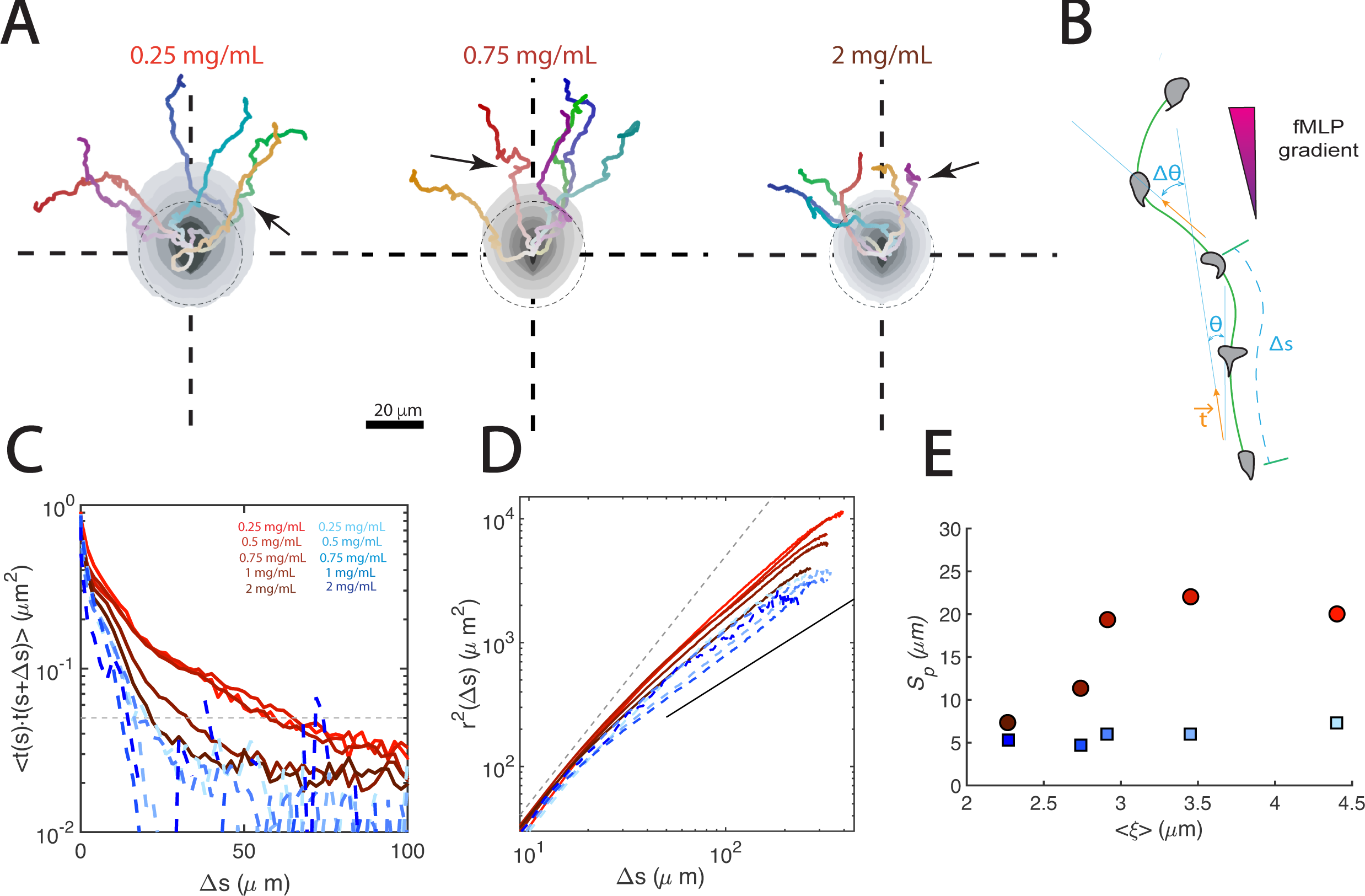
Persistence of Cell Trajectories vs. Matrix Density. **A)** Probability density maps of cell coordinates (gray contours), and representative cell trajectories for non-restrictive ([col] = 0.25 mg/ml), intermediate ([col] = 0.75 mg/ml), and restrictive ([col] = 2 mg/ml) matrices. Arrows indicate representative turns. Time progression along each trajectory is represented by changes from white to saturated colors. The dashed circles represent unbiased random motion **B)** Schematic of a cell trajectory, indicating a segment of length Δs, along which the tangent vector 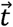 changes its orientation *θ* with respect to the gradient by *Δθ*. **C)** Tangent vector autocorrelations as a function of distance traveled, *C*_*tt*_ (Δ*s*), for no fMLP (dashed blue lines) and fMLP (red solid lines) conditions, in matrices of varying density. The dashed line represents the cutoff *C*_*tt*_= 0.05 used to determine the persistence length of the trajectories (*S*_*p*_). **D)** Mean squared displacements vs. distance separation along cell trajectories, *r*^2^ (Δ*s*). The dashed and solid lines represent *r*^2^∼Δ*s*^2^ and *r*^2^∼Δ*s* respectively. **E)** Persistence length of the cell trajectories, *S*_*p*_, as a function of matrix mean pore size, ⟨*ξ*⟩, for no fMLP (blue squares) and fMLP (red circles) conditions.

Next, we determined the mean squared displacements (MSD) of the cells’ trajectories. The MSD versus time, *i*.*e*., 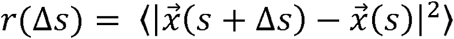, have been widely used to quantify 2-D and 3-D cell migration by exploiting the analogy between these processes and a random walk [22, 24, 44-46]. In particular, the logarithmic slope of the MSD plot is often used to assess whether migration is random or directed. In the present study, we considered the MSD as a function of distance traveled, *i*.*e*., 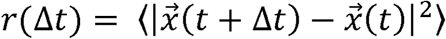 because this function characterizes the geometry of the cell trajectories independent of cell speed, whereas *r*(Δ*t*) does not. This consideration is relevant because cell speed can depend on [col] and the local curvature of the trajectory, as will be shown below. The rate of increase of the MSD with Δ*s* depends on whether cells keep memory of their direction of translocation over a distance equal to Δ*s*. Note that since the MSD can be expressed as

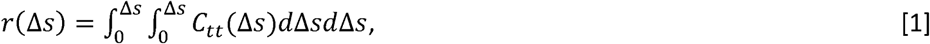

the MSD are tightly linked to the autocorrelation of the direction of cell migration. The MSD of chemotaxing and randomly moving cells were plotted in Figure 3D for various collagen concentrations. For short travel distances, *r*(Δ*s*) ∼ Δ*s*^2^ in all conditions tested, a hallmark of directed motion that is consistent with the *C*_*tt*_ ≈ 1values that are shown in Figure 3C. Subsequently, the MSD tapered towards *r*(Δ*s*)∼Δ*s* as Δ*s* increased and the cells began to add changes of direction (turning events) along their paths as *C*_*tt*_ progressively dropped.

The rate of decay of *C*_*tt*_ governs the transition between the linear and quadratic regimes of the MSD according to Equation 1, and is related to the persistence length of the trajectories, *S*_*p*_ [47]. Thus, we estimated *S*_*p*_ using an exponential decay model for *C*_*tt*_ (see Methods for details) and plotted the results in Figure 3E. These data show that the persistence of the paths of neutrophils undergoing directed migration through 3-D matrices plateaued at s_*P*_ ≈20 *μm* for matrices with ⟨*ξ*⟩ ≳ 3 *μm*, and then fell sharply to reach s_*P*_ ≈ 7*μm* for ⟨*ξ*⟩ < 3 *μm*. In contrast, randomly moving neutrophils had trajectories with *s*_*P*_ ≈5 *μm* regardless of ⟨*ξ*⟩. Together with the matrix deformation measurements presented in Figure 2, these results suggest that neutrophils undergoing chemotaxis perform more frequent turns in restrictive matrices with narrower pores that cannot be easily deformed. In our most restrictive matrices, these turns became almost as frequent as those performed by randomly moving cells.

Previous studies have uncovered key features of 3-D random cancer cell migration by characterizing the probability distribution of turning angles in cell paths [24]. Thus, we determined the distribution of *Δθ* vs. *Δs* for the chemotactic neutrophil trajectories tracked in our 3-D experiments, *i*.*e*., *p*(*Δθ*|*Δs*). Figures 4A-B display contour maps of *p*(*Δθ*|*Δs*) in 0.5 mg/ml and 2 mg/ml collagen matrices. They show that Δθ is narrowly distributed around *Δθ = 0* for small *Δs*, implying that the neutrophils followed relatively straight paths for short spatial separations. With increasing *Δs*, the distributions become progressively more uniform as the neutrophils implement changes of direction to navigate the matrix and their paths lose directional persistence. As expected from the *C*_*tt*_ and MSD data described above, the loss of persistence took place at shorter distances with increasing [col]. However, it is worth noting that, while the distribution of *Δθ* became quasi uniform at *Δs* ∼ *20 μm* for randomly moving neutrophils (insets in Figure 4A-B), chemotaxing neutrophils exhibited directional memory after traveling long distances, as denoted by *p*(*Δθ*|*Δs* > *100 μm*) which still maintains a peak at *Δθ=0*. This persistence was also observed in the most restrictive matrices, despite the cells turning more often in these matrices.

**Figure 4.**
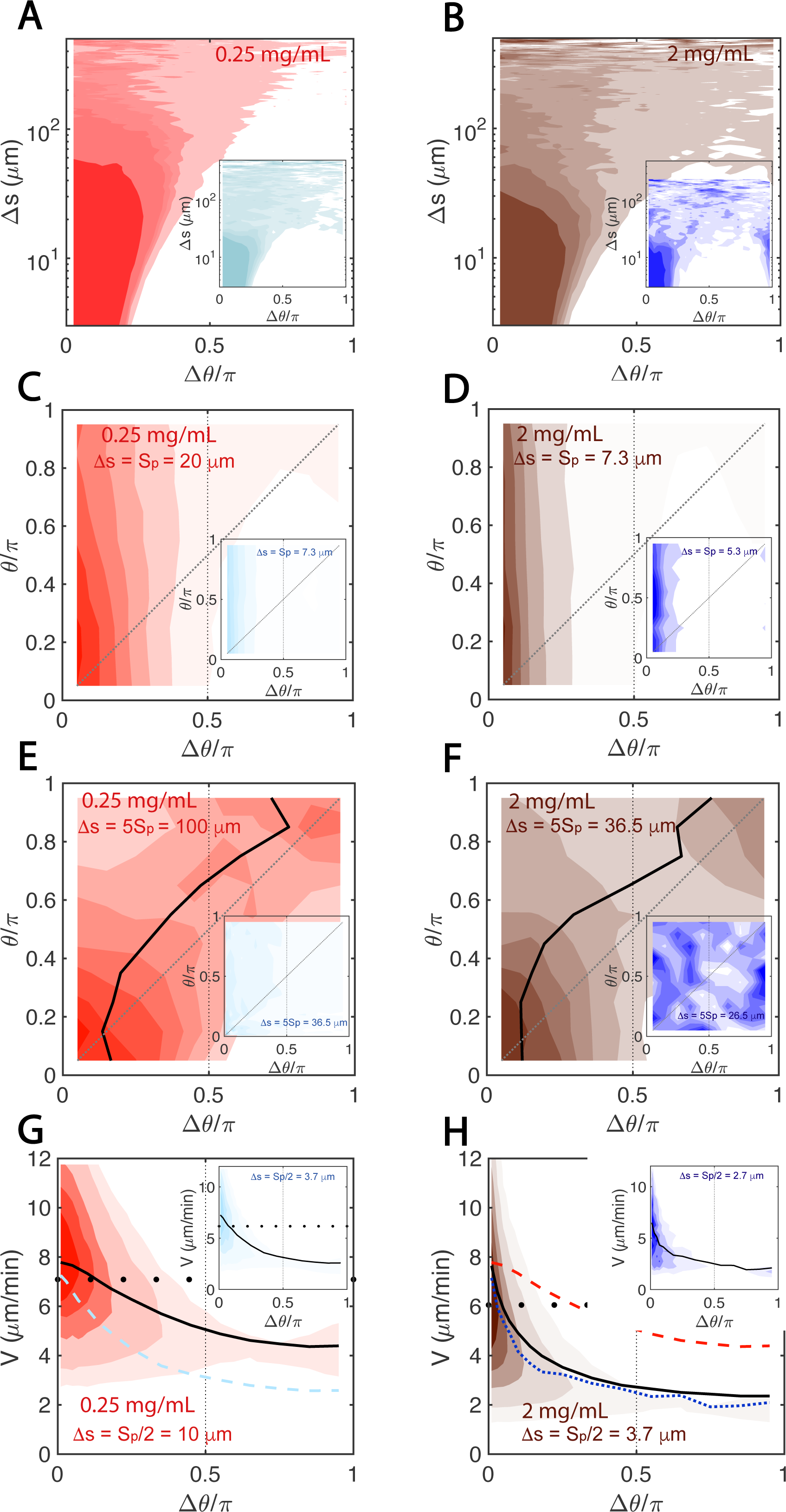
Statistics of Turning angles and Cell Speeds vs. Matrix Density. **A-B)** Maps of probability distribution of turning angles Δ *θ* conditional to distance separation Δs along the trajectory, p (Δ *θ* |Δ*s*). C-D) Maps of probability distribution of Δ *θ* conditional to the orientation e with respect to the gradient, for a distance separation equal to the persistence length, p (Δ*θ* | *θ*, Δ*s* = *s*_*p*_). **E-F)** Maps of *p* (Δ*θ* | *θ*, Δ*s* = 5 *s*_*p*_). **G-H)** Maps of joint probability distribution of turning angle and cell speed, for a distance separation Δ*s* = *s*_*p*_/2, *i*.*e*., *p* (Δ *θ, v*, Δ*s* = *s*_*p*_/2). The maps in panels **A, C, E**, and **G** come from 0.5 mg/ml matrices, whereas those in panels **B, D, F**, and **H** come from 2mg/ml. In each panels the main probability map comes from fMLP conditions, while the inset comes from randomly moving cells in the absence of fMLP. The dotted and solid lines in panels **C-F** indicate respectively *θ* = Δ *θ* and *θ* = Δ *θ* _*m*_ (*θ*) where Δ *θ* _*m*_ is the mode of the Δ *θ*-distribution. The solid and dotted lines in panels **G-H** represent respectively the mean cell speeds of chemotaxing and randomly moving cells vs. turning angle for 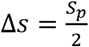, *i*.*e*., *v* (Δ*θ*, Δ*s* = *s*_*p*_ /2). The solid circles in panels **G-H** indicate the total mean speed ⟨*v*⟩ for each matrix (shown in Figure 1F). In panel **H**, the dashed red line represents the mean speed of chemotaxing cells in the 0.5 mg/ml matrix.

### 3.5. Chemotaxing neutrophils integrate short-range displacements dictated by diverse 3-D microenvironments to achieve long-range directional bias

The data presented above suggest that the persistence length of neutrophils migrating in 3-D is dictated by small-scale features of the matrix such as pore size. To investigate how these cells balance the short-range persistence imposed by their microenvironment with the long-range directional bias necessary for chemotaxis, we calculated the probability distribution of their turn angles conditional to their prior orientation with respect to the gradient and distance separation, i.e., *p*(*Δθ /θ,Δs*) where 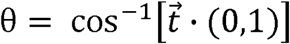 (see diagram in Figure 3B). Figures 4C-D show maps of *p*(*Δθ /θ,Δs*) for a fixed distance separation *Δs* = *S*_*p*_ for [col] = 0.5 and 2 mg/ml. Because of the definition of *S*_*p*_, this distance separation corresponds to one turn on average. In both matrices, *p*(*Δθ /θ,Δs=S*_*p*_) is independent of *θ*, indicating that neutrophils do not consider the direction of the chemoattractant gradient when performing a single turn. In fact, the angular distributions of chemotaxing neutrophils were similar to those for randomly moving ones at this distance separation (see insets in Figures 4C-D).

Next, we examined the conditional *Δθ* −distributions of turning angles for segments of trajectory concatenating multiple turns, i.e., *p*(*Δθ /θ,Δs=5 S*_*p*_) (Figures 4E-F). Regardless of [col], these distributions peaked near Δθ=θ, in evident contrast with the distributions obtained from randomly moving cells (Figure 3E-F, inset), which were roughly uniform and independent of θ. Overall, our data are consistent with a model in which 3-D migrating neutrophils implement turns in their trajectories to circumvent impassable sections of the matrix. These short-range turns are dictated by the local microenvironment and result in misalignments with respect to the gradient that are not immediately corrected. Instead, it takes several consecutive turns for neutrophils to realign themselves with the gradient in both non-restrictive and restrictive matrices. Section 3.9 examines this realignment process in more detail.

### 3.6. Neutrophils migrating in 3-D decrease their speed when turning

To study how turning events affected the speed of cell migration, we generated probability distributions of cell velocity and *Δθ* as a function of the distance traveled in our different matrices, *i*.*e*., *p*(*Δθ,v*|*Δs*). We observed a clear dependence of *v* on *Δθ* when we considered a segment with approximately one turn by choosing *Δs* comparable to *S*_*p*_. To illustrate this dependence, we plotted 2-D probability density maps of *p*(*Δθ,v*|*Δs*) for fixed *Δs = S*_*p*_*/2* (Figures 4G-H). These data revealed that neutrophil speeds were reduced significantly while turning, and that this reduction was greater for sharper turns (i.e., higher *Δθ*). When [col] increased from 0.5 to 2 mg/ml, cell speed decreased further (compare red dashed and black solid lines in Figure 4H), and this decrease was more pronounced when the cells were turning. Given that cells turned more often as [col] increased, the difference between the speed of straight motion (v(*Δθ=0*)) and the mean cell speed (solid circles in Figures 4G-H) was larger in high-[col] matrices than in low-[col] ones.

### 3.7. Short-range persistence impedes long-range directional bias in restrictive matrices

After finding that neutrophils increase their frequency of turning in restrictive matrices, thereby decreasing their migratory persistence, we examined how perturbations leading to increased persistence would affect 3-D cell migration in different microenvironments. To this end, we treated cells with the Arp2/3 inhibitor ck666, which restricts the ability of the Arp2/3 complex to polymerize branched actin filament networks and form lamellipodial protrusions [48, 49]. Previous studies have postulated that Arp2/3-mediated lamellipodia formation is crucial for exploring the local microenvironment during 3-D migration and finding the best paths (pores) to migrate through [11, 17]. These studies have also shown that the inhibition of the Arp2/3 complex leads to cell trajectories that are more persistent in space.

We first examined the probability distributions of cell positions after ck666 treatment and found a consistent trend of decreased travelled distances and straighter trajectories compared to untreated cells, even for high [col] (compare Figure 5A and Figure 3A). Consistently, the turning angles of ck666-treated cells were narrowly distributed near *Δθ=0* for *Δ*s<100 μm regardless of [col] (compare Figure 5A and Figures 4A-B). Relative to untreated cells, the MSDs vs Δ*s* of the ck666-treated cells remained relatively close to *r*(Δ*s*) ∼ Δ*s*^2^ (Figure 5B), and *C*_*tt*_ decayed more slowly for these cells than for untreated cells (Figure 5C). Consequently, the persistence distance of the migrating ck666-treated cells was generally higher than that of untreated cells (Figure 5D). Similar to untreated cells, we observed a transition towards shorter *S*_*p*_ with decreasing *ξ*. Furthermore, this transition seemed to occur for larger *ξ* than in untreated cells, suggesting that the migration of Arp2/3-inhibited cells was disrupted by less restrictive matrices than the migration of untreated cells. However, ck666-treated cells were unable to reduce their persistence as much as untreated cells to navigate matrices with small pore sizes. For instance, *Sp*(*ξ=2*.*3 μm*) was twice as long for ck666-treated than for untreated cells (14 vs 7 *μm*, see Figure 5D).

**Figure 5.**
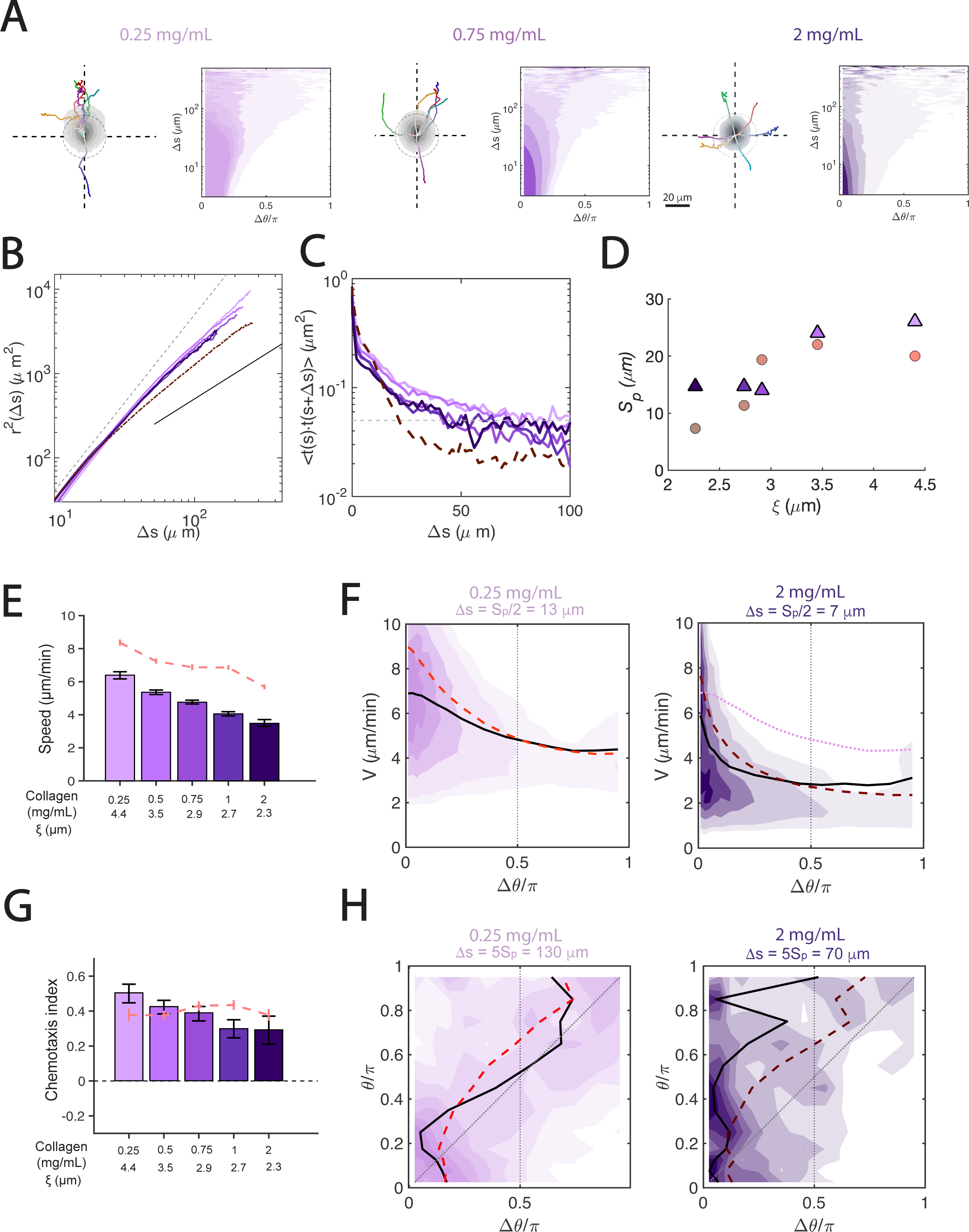
Migration Statistics for Arp2/3-inhibited cells. **A)** Probability density maps of cell coordinates and representative cell trajectories together with maps of *p* (Δ*θ* |Δ*s*) for non-restrictive ([col] = 0.25 mg/ml), intermediate ([col] = 0.75 mg/ml), and restrictive ([col] = 2 mg/ml) matrices. The data come from chemotaxing cells treated with the Arp2/3 inhibitor ck666. Time progression along each trajectory is represented by changes from white to saturated colors. The dashed circles represent unbiased random motion. **B)** Mean squared displacements of ck666-treated cells in matrices of varying density, *r*^2^ (Δ*s*). The dashed and solid lines represent *r*^2^∼Δ*s*^2^ and *r*^2^∼Δ*s* respectively. **C)** *C*_*tt*_ (Δ*s*) of ck666-treated cells in matrices of varying collagen density. The grey dashed line represents the cutoff a = 0.05 used to determine *S*_*p*_. In **B** and **C**, the dashed dark red lines come from untreated cells in the 2 mg/ml matrix. **D)** Persistence length of the cell trajectories, *S*_*p*_, as a function of matrix mean pore size, ⟨*ξ*⟩, for ck666-treated cells (purple triangles) and untreated cells (red circles). **E)** Mean speeds <*v*> of ck666-treated cells vs. matrix density. The red dashed line represents <*v*> from untreated cells. Error bars represent 95% confidence interval. **F)** Maps of *p* (Δ*θ*, v, Δ*s* = *s*_*p*_/2) for ck666-treated cells in non-restrictive ([col] = 0.25 mg/ml, left panel) and restrictive ([col] = 2 mg/ml, right panel) matrices. The solid and dashed lines represent the mean cell speed *v* (Δ*θ*, Δ*s* = *s*_*p*_/2) for ck666-treated cells and untreated cells, respectively. The dotted line in the right panel represents *v* (Δ*θ*, Δ*s* = *s*_*p*_/2) for ck666-treated cells in the 0.25 mg/ml matrix. **G**) Chemotactic index (CI) of ck666-treated cells vs. matrix density. The red dashed line represents *CI* from untreated cells. Error bars represent 95% confidence interval. **H)** Maps of *p* (Δ*θ*| *θ*, Δ*s* = 5 *s*_*p*_) for ck666-treated cells in sparse (0.25 mg/ml, left panel) and dense (2 mg/ml, right panel) matrices. The dotted and solid lines indicate respectively *θ* = Δ*θ* and *θ*= Δ*θ*_*m*_(*θ*). The dashed red lines correspond to Δ*θ*_*m*_ (*θ*) from untreated cells.

On average, ck666 treatment caused a decrease in neutrophil speed, and this decrease was more accentuated for high [col] (Figure 5E). A more detailed analysis of neutrophil trajectories revealed that, similar to untreated neutrophils, the speed of ck666-treated cells dropped mildly with *Δθ* for [col] = 0.25-mg/ml (Figure 5F, left panel) and sharply for [col] = 2.0-mg/ml (Figure 5F, right panel). Of note, the largest differences in speed between untreated cells and ck666-treated cells occurred at low turning angles in both matrices (compare black lines and red dashed lines in Figure 5F), implying that Arp2/3 inhibition not only impaired path finding, but also slowed down straight motion.

Next, we asked how the increased short-range persistence (i.e., higher *S*_*p*_) caused by Arp2/3 inhibition affected chemotactic efficiency, as measured by CI. Our results (Figure 5G) indicated that ck666-treated cells had a higher CI than untreated cells for low [col]. However, the CI of ck666-treated cells decreased with [col], falling below the CI of untreated cells in our three highest-[col] matrices. In an effort to understand these trends, we analyzed the *p*(*Δθ /θ,Δs = 5S*_*p*_) for ck666-treated cells in the 0.5 and 2 mg/ml matrices (Figure 5H). Our analysis showed that, despite their decreased rate of turning (i.e., increased *S*_*p*_), the Δθ −distributions peaked near *Δθ=θ* in the 0.25 mg/ml matrix (Figure 4H, left panel). Thus, ck666-treated cells were able to undergo sufficient turning to maintain directional bias towards the chemoattractant source in non-restrictive matrices, similar to untreated cells. However, the *Δθ* −distributions from ck666-treated cells were strikingly different in restrictive, high-[col] matrices (Figure 5H, right panel). In particular, they became wider with an increasing misalignment angle *θ* but peaked near *Δθ=0* regardless of *θ*. Data from other values of *Δs* yielded similar results. These findings suggest that the ck666-treated cells underwent decreased biased turning and were not able to correct their misalignment with respect to the chemotaxis axis in restrictive matrices.

### 3.8. Myosin-II contractility is not crucial for turning but facilitates fast 3-D neutrophil migration

Previous studies have shown that myosin-II-mediated rear contractility is an important mechanism in 3-D cell migration [50]. Inhibition of myosin-II significantly decreases the ability of neutrophils to squeeze through narrow constrictions [16]. Given that our data for both untreated and ck666-treated cells suggested an important role for turning in maintaining the long-range directional bias required for chemotaxis, we examined whether myosin-II contractility was necessary for turning. To investigate this, we treated cells with the non-muscle myosin-II inhibitor blebbistatin, which disrupts myosin-II mediated cell contractility [51, 52].

The distributions of cell positions after blebbistatin treatment (Figure 6A) suggested decreased traveled distances when compared to untreated cells (Figure 3A) for all collagen concentrations; however, a comparison of the 6 longest cell trajectories did not reveal noticeable differences in turning events. As with untreated cells, the turning angles for blebbistatin-treated cells were narrowly distributed around *Δθ=0* for small *Δ*s and progressively became more uniform with increasing *Δ*s (Figure 6A). The MSDs vs. Δ*s* results for blebbistatin-treated cells were also similar to those for untreated cells in that *r*(Δ*s*) ∼ Δ*S*^2^ for small *Δ*s and tapered to *r*(Δ*S*)∼Δ*S* as Δ*S* increased (Figure 5B). However, careful analysis of the persistence of blebbistatin-treated cells from their *C*_*tt*_ (Δ*S*) (Figure 6C) revealed that decreased contractility made cells more sensitive to their microenvironment, as the transition to an increased rate of turning occurred for larger ξand lower [col] than in untreated cells (Figure 6D).

**Figure 6.**
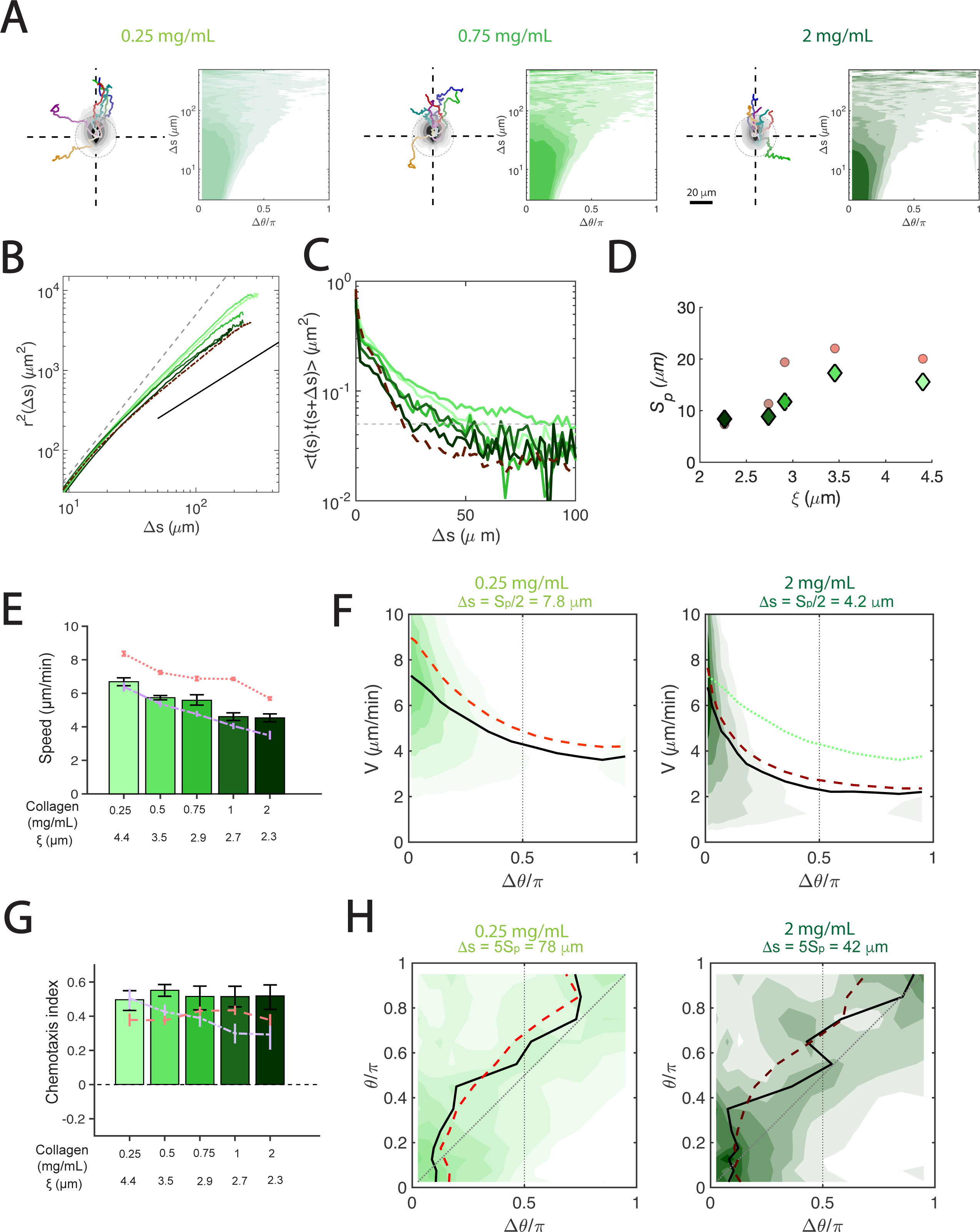
Migration Statistics for myosin II-inhibited cells. **A)** Probability density maps of cell coordinates and representative cell trajectories together with maps of p (Δ*θ*|Δs) for non-restrictive ([col] = 0.25 mg/ml), intermediate ([col] = 0.75 mg/ml), and restrictive ([col] = 2 mg/ml) matrices. The data come from chemotaxing cells treated with the myosin II inhibitor blebbistatin. Time progression along each trajectory is represented by changes from white to saturated colors. The dashed circles represent unbiased random motion. **B)** Mean squared displacements of blebbistatin-treated cells in matrices of varying collagen density, *r*^2^ (Δ*s*). The dashed and solid lines represent *r*^2^∼Δ*s*^2^ and *r*^2^∼Δ*s* respectively. **C)** *C*_*tt*_ (Δ*s*) of blebbistatin-treated cells in matrices of varying density. The grey dashed line represents the cutoff *C*_*tt*_= 0.05 used to determine *S*_*p*_. In **B** and **C**, the dashed dark red lines come from untreated cells in the 2 mg/ml matrix. D) Persistence length of the cell trajectories, *S*_*p*_, as a function of matrix mean pore size, ⟨*ξ*⟩, for blebbistatin-treated cells (green diamonds) and untreated cells (red circles). **E)** Mean speeds <*v*> of blebbistatin-treated cells vs. matrix density. The red and purple dashed lines represent <*v*> from untreated and ck666-treated cells, respectively. Error bars represent 95% confidence interval. **F)** Maps of *p* (Δ*θ*, v, Δ*s* = *s*_*p*_/2) for blebbistatin-treated cells in sparse (0.25 mg/ml, left panel) and dense (2 mg/ml, right panel) matrices. The solid and dashed lines represent the mean cell speed *v* (Δ*θ*, Δ*s* = *s*_*p*_/2) for blebbistatin-treated cells and untreated cells, respectively. The dotted line in the right panel represents *v* (Δ*θ*, Δ*s* = *s*_*p*_/2) for blebbistatin-treated cells in the 0.25 mg/ml matrix. **G)** Chemotactic index (CI) of blebbistatin-treated cells vs. matrix density. The red and purple dashed lines represent *CI* from untreated cells. Error bars represent 95% confidence interval. **H)** Maps of *p* Δ*θ* |*θ*, Δ*s* = 5 *s*_*p*_ for blebbistatin-treated cells in sparse (0.25 mg/ml, left panel) and dense (2 mg/ml, right panel) matrices. The dotted and solid lines indicate respectively *θ* = Δ*θ* and *θ* = Δ*θ* _*m*_ *θ*). The dashed red lines correspond to Δ*θ* _*m*_ *θ*) from untreated cells.

Blebbistatin treatment resulted in decreased average neutrophil speeds that continued to decrease with [col] (Figure 6E). This drop in speed was nearly constant across all turning angles (Figure 6F). The CI of blebbistatin-treated cells did not vary with [col] and was slightly higher than that for untreated cells (Figure 6G). Consistent with the CI data, the long-range (i.e., Δs = *5 S*_*p*_) *Δθ*−distributions conditional to *θ* did not vary qualitatively after blebbistatin treatment in both non-restrictive and restrictive matrices (Figure 6H). Together, these results suggest that myosin-II contractility is important for fast 3-D neutrophil migration and affects the transition to decreased short-range persistence as collagen concentration increases. However, neutrophils are able to navigate restrictive matrices by coordinating consecutive turning events to maintain long-range directional bias with or without fully active myosin-II contractility.

### 3.9. Directional bias depends on Arp2/3 and requires on average three consecutive turning events, regardless of matrix density and myosin-II contractility

Neutrophils chemotaxing on 2-D surfaces have been shown to immediately correct their misalignments with respect to a chemotactic gradient in one turn [53]. However, our data suggests that in 3-D matrices, these cells needed to coordinate several turns over a directional bias distance to achieve the same objective (Figure 4). We denoted this distance *S*_*θ*_ (see diagram in Figure 7A) and estimated it in two steps. First, we obtained *p*(*Δθ,v*|*Δs*) as described above and calculated the difference between the most likely turning angle in this distribution, Δ *θ*_*m*_(*θ*,Δ*s*), and a full directional correction (i.e., Δ *θ*_*m*_(*θ*,Δ*s*) = *θ*) for each value of Δ*s*,

**Figure 7.**
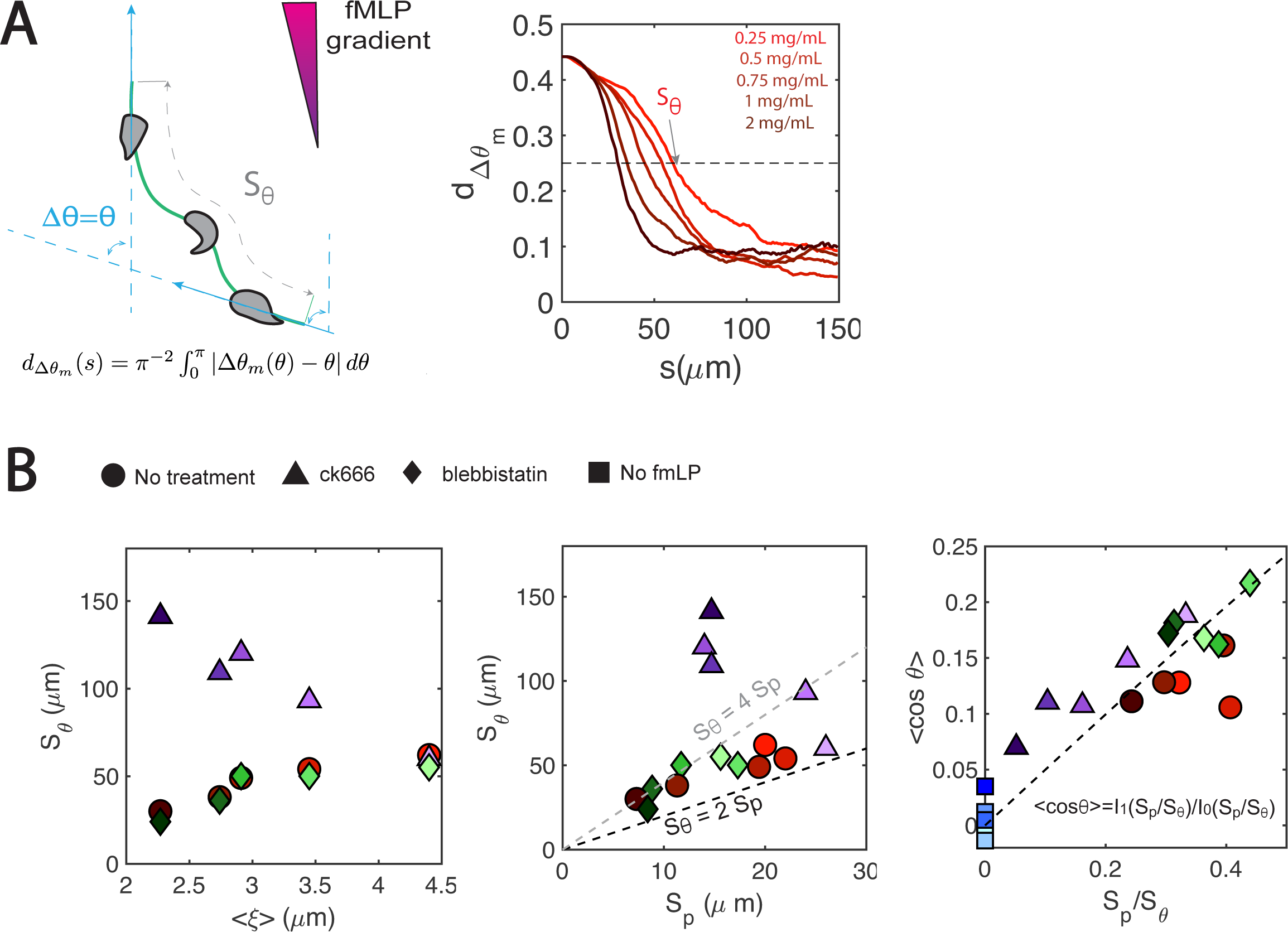
Realignment with the chemoattractant gradient. **A)**<u>Left:</u> Schematic indicating the bias lengthscale, s_θ_, defined as the distance that cells need to correct an angular misalignment e with respect to the chemoattractant gradient. <u>Right:</u> Metric 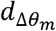 (see left panel for equation) that quantifies how close cells are to a total alignment correction Δθ = θ as a function of distance separation Δs. The data comes from untreated chemotaxing cells in matrices of different densities. The grey dashed line indicates the threshold used to determine s_*θ*_. **B)** <u>Left</u>: *s*_*θ*_vs.⟨*ξ*⟩ for different treatments and matrix densities. <u>Right:</u> *s*_*θ*_ vs. *s*_*p*_ for different treatments and matrix densities. The dashed dark and light lines represent *s*_θ_ = 2*s*_*p*_ and *s*_θ_ = 4*s*_*p*_ respectively. Untreated cells, ck666-treated cells and blebbistatin-treated cells are represented by red circles, purple triangles and green diamonds respectively, and their color indicates collagen density (0.25, 0.5, 0.75, 1 and 2 mg/ml from light to dark).

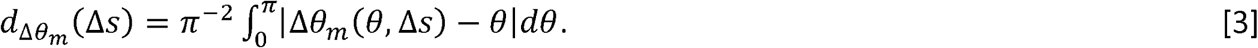

It is straightforward to show that this definition yields 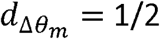 for a narrow (e.g., Dirac delta) distribution with Δ *θ*_*m*_= 0. Consistently, we found that 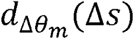 decreases with Δ*s* from its value 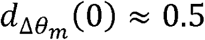 at the origin (Figure 7A, right panel). The second step to determine *S*_*θ*_ was to use the intersection between the 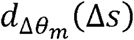 curve and the threshold value 1/4 to estimate *S*_*θ*_ for each collagen concentration, both for untreated and pharmacologically manipulated cells. We chose the 1/4 threshold because 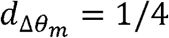 for a uniform distribution with Δ *θ*_*m*_= π/2.

The results of this analysis (Figure 7B, left) indicated that untreated cells and blebbistatin-treated cells were able to correct misalignments with respect to the gradient over a distance *S*_*θ*_ ≈50 *μm* in non-restrictive, low-[col] matrices with ⟨*v*⟩ > 3 *μm*. However, this distance decreased to *S*_*θ*_ ≈30 μm in restrictive, high-[col] matrices with ⟨*ξ*⟩< 3 *μm*. To investigate this seemingly counter-intuitive result that cells can restore their chemotactic compass over a shorter distance in more restrictive matrices, we plotted *S*_*θ*_ vs. the persistence distance *S*_*p*_ (Figure 7B, center). Interestingly, we found that *S*_*θ*_ spanned a relatively narrow range when scaled with *S*_*p*_ [*S*_*θ*_ ≈(2 − 4) *S*_*p*_]. This result implies that it takes neutrophils between two and four turns on average to realign their motion with the chemoattractant gradient. Furthermore, since cells turn more often in more restrictive matrices, they need a shorter physical distance to realign. Our data suggest that collagen concentration or myosin-II contractility did not seem to affect this result, but ck666-inhibition of the Arp2/3 complex drastically increased *S*_*θ*_ (Figure 7B, center). Thus, ck666-treated cells that performed turns to avoid obstacles in their restrictive microenvironments had a reduced ability to control the direction of turning to remain aligned with the chemotaxis gradient.

Given that *S*_*P*_ and *S*_*θ*_. can be loosely interpreted as the frequencies per unit distance of random and biased turns along a cell’s trajectory, we hypothesized that the chemotactic efficiency of the cells would depend on the ratio *S*_*P*_/*S*_*θ*_. To test this hypothesis, we plotted the average of cos *θ* vs. *S*_*P*_/*S*_*θ*_ for all our experimental conditions (Figure 7B, right). These data confirm that cell motion remained more aligned with the chemoattractant gradient for larger values of *S*_*p*_ /*S* _*θ*_. In addition, we built mathematical models of biased random walks with persistence distance *S*_*p*_ and bias lengthscale *S*_*θ*_ arising from an angular controller

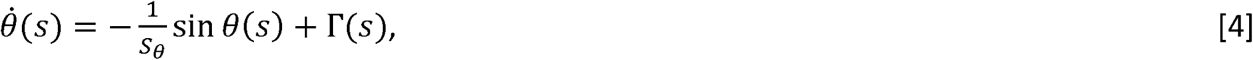

where Γ (*s*) is a white noise source with 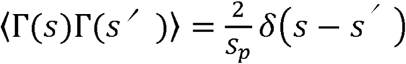 (see Supplementary Information and refs. [54]. The simplest version of these models, which assumes constant cell speed independent of *θ*, is particularly interesting because it has an exact analytical solution for the average of cos *θ* that does not have any adjustable parameter, *i*.*e* 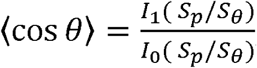 where *I*_*1*_ and *I*_*0*_ are modified Bessel functions of the first kind. This model prediction agrees well with our experimental data (Figure 7B, right).

### 3.10 Cell-generated mechanical forces affect the rate of turning and speed of neutrophils migrating in 3-D, but not their chemotactic efficiency

We examined to what extent neutrophils relied on mechanical forces to migrate in 3-D environments by quantifying the matrix deformations caused by untreated, ck666-treated and blebbistatin-treated cells. In our 0.25 mg/ml matrices, both treatments caused a notable decrease in 3-D matrix deformations (Figure 8A-C) that was statistically significant in the case of blebbistatin treatment as measured by the 3-D contraction moment (|*tr* (***M***)| defined in eq. 2, Figure 8D); however, the treated cells were still able to appreciably deform their microenvironment. Given that myosin-II inhibition caused a marked decrease in *S*_*p*_ for intermediate [col] (*i*.*e*., 0.75 mg/ml, see Figure 5D), we measured the contraction moments created by blebbistatin-treated cells in these matrices (Figure 8E), and found that |*tr* (***M***)| was significantly reduced after treatment. The contractile moments in our most restrictive matrices were about 10 times lower than in non-restrictive matrices, and also decreased after inhibiting myosin-II contractility or Arp2/3-mediated branched actin polymerization (Figure 8F).

**Figure 8.**
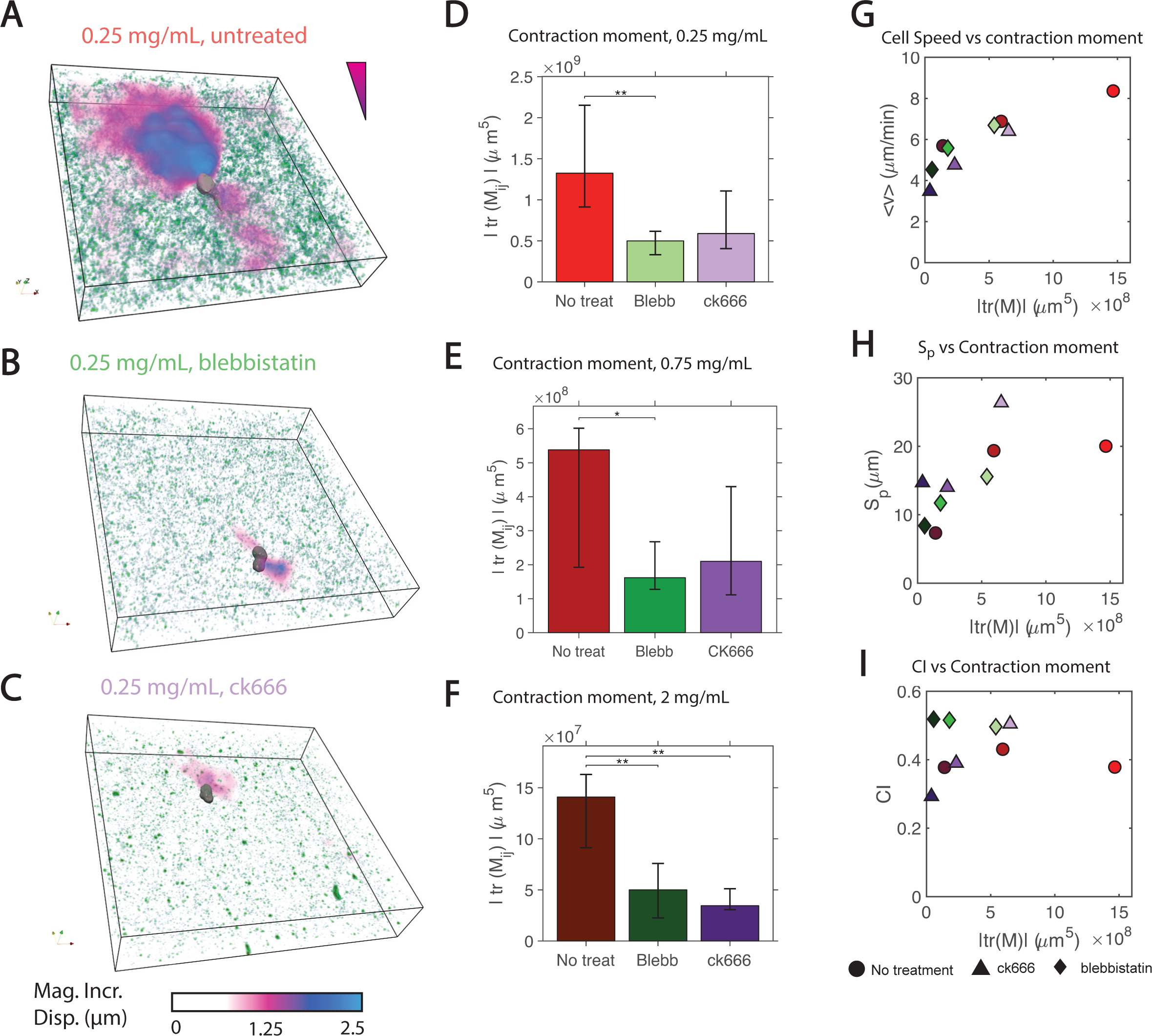
Relationship between cell-generated matrix deformations and cell migration parameters. **A-C)** Representative reconstruction of cells (grey), beads (green), and incremental 3-D matrix deformation (magenta – blue colormap) during migration in a sparse (0.25 mg/ml) collagen matrix for no treatment **(A)**, blebbistatin treatment **(B)**, and ck666 treatment **(C). D-F)** Mean magnitude of 3-D contractile moment, |*tr* (***M***)|, for different treatments in sparse (0.25 mg/ml, panel **D)**, intermediate (0.75 mg/ml, panel **E)**, and dense (2 mg/ml, panel F) matrices. Error bars represent 95% confidence intervals of the mean. * and ** respectively indicate p<0.05 and p<0.01 according to the Mann-Whitney U test. **G)** Mean cell speed <*v*> vs. |*tr* (***M***)| for different treatments and matrix densities. **H)** Persistence length of cell trajectories, *S*_*p*_, vs. |*tr* (***M***)| for different treatments and matrix densities. **I)** Chemotaxis index *CI* vs. |*tr* (***M***)| for different treatments and matrix densities.

Since blebbistatin- and ck666-treated cells had lower migration speeds than untreated cells, we examined the relationship between |*tr* (***M***)| and ⟨*v*⟩ (Figure 8G). This analysis revealed that |*tr* (***M***)| and ⟨*v*⟩ collapsed on a single curve regardless of treatment and [col], suggesting that the ability of 3-D migrating neutrophils to mechanically remodel their environment is a key factor governing their speed. Given that cell speed decreased when cells were turning (Figures 4G-H, 5F and 6F), we plotted *S*_*p*_ vs. |*tr* (***M***)| (Figure 8H) and found that, for both treatments and control conditions, *S*_*p*_ decreased when |*tr* (***M***)| fell below ∼ 5×10^3^ *μm*^2^ regardless of [col]. Thus, neutrophils may transition to a migratory strategy with an increased rate of turning when they become unable to sufficiently deform their surrounding matrix. Finally, we plotted the chemotaxis index vs. |*tr* (***M***)| (Figure 8I) and found that this index did not depend on |*tr* (***M***)|except when Arp2/3 was inhibited.

## Discussion

The present study investigated the ability of neutrophils to migrate up a chemoattractant gradient in 3-D environments with different degrees of complexity. We hypothesized that turning events would be a key factor for quick migration towards a chemoattractant source by allowing cells to circumvent obstacles in the matrix (e.g., narrow pores) and quickly resume their motion towards the chemoattractant. We tested these hypotheses using a custom-made 3-D collagen matrix chemotaxis chamber, label-free automated cell-tracking, and statistical analyses to quantify the migratory trajectories of neutrophil-like HL-60 cells and the matrix deformation these cells exert during migration.

Previous studies used time-dependent kinematic analyses of leukocyte trajectories in 3-D matrices to find that cell speed and persistence decrease with decreasing matrix pore size [11, 21, 23]. Although more scarce, there are previous reports of 3-D matrix deformations caused by migrating leukocytes [55]. In the present study, we analyzed cell trajectories based on distance traveled rather than time elapsed during migration. This interpretation allowed us to study the geometry of cell trajectories and decouple it from the speed of cell migration, which varies along the trajectories as the cells navigate different matrix sections. Furthermore, our label-free automated cell-tracking made it possible to analyze over 20,000 migrating cells, yielding highly detailed, multi-parametric statistics of the migratory process. Together with our 3-D measurements of matrix deformation, these data have revealed mechanistic interplays between cell contractility and migratory parameters that had not been quantified before.

We first characterized how increasing matrix collagen concentration ([col]) caused a decrease in matrix pore (ξ) size and heterogeneity, leading to a stark reduction in available pores larger than the cell nucleus in our most restrictive matrices. We found that the decreasing ξ slowed cells down but did not diminish their ability to migrate directionally up the chemoattractant gradient, consistent with previous studies involving other leukocyte subtypes [21, 23]. Our measurements of 3-D matrix deformations indicated that neutrophils could transiently deform low-[col] matrices using physical forces; however, they became unable to appreciably deform high-[col] matrices. The cells adapted to increased matrix rigidity in high-[col] matrices by turning more often, which caused a marked decrease in the spatial persistence of the direction of motion. This transition was triggered by the cells’ inability to physically remodel their microenvironment, which was confirmed by pharmacological inhibition of contractile forces mediated by myosin-II, or protrusive forces exerted by Arp2/3 – mediated dendritic actin polymerization.

The physical properties of different types of tissue, in which the extracellular matrix is a significant component, can vary greatly across different locations in the body [56]. However, during immune responses, this variation does not impede neutrophils from migrating towards sites that initiate inflammatory responses [8]. The extracellular matrices in our study contained average pore sizes well within physiological ranges [57]. Therefore, the versatility in migration strategies that we found *in vitro* has implications for understanding the ability of leukocytes to migrate across a range of environments during innate immune responses [9]. This adaptability is crucial for effective chemotaxis because, unlike cancer cells, neutrophils do not heavily rely on proteolytic matrix degradation when facing restrictive 3-D environments [21, 58]. Cancer cells can transition to proteolytic matrix degradation when presented with rigid microenvironments that they cannot deform by means of physical forces [59]. The ensuing permanent matrix remodeling has been shown to cause migratory statistics consistent with an anisotropic random walk, thus ensuring a long-range directional bias [24]. In contrast, 3-D migrating neutrophils adapt to non-compliant microenvironments by turning more often; thus, long-range directional bias is achieved by a different means.

In our experiments, the short-range statistics of 3-D neutrophil migration became progressively disturbed with decreasing ξ, becoming similar to those of randomly moving cells. However, neutrophils conserved their chemotactic efficiency, measured as the ratio of distance traveled towards the chemoattractant to total distance traveled, even in the most restrictive matrices. This result hints at a tradeoff between short-range and long-range goals in the directed 3-D migration of neutrophils. A key factor in this tradeoff is the ability to correct the misalignments with respect to the gradient that arises when circumventing impassable sections of the matrix. Our data show that cells barely consider their orientation with respect to the gradient when performing one single turn, but they progressively integrate this information into subsequent turns and eventually realign their motion with the gradient. We found that a biased random walk model [54] with no adjustable parameters reproduced the evolution of cell motion orientation along the cell trajectories. In the biased random walk model, cells steer towards the chemoattractant source over a bias distance, *s*_*θ*_, which dictates the chemotactic efficiency in competition with the persistence of cell migration, *s*_*p*_; *i*.*e*., cells with higher values of *s*_*p*_ /*s*_*θ*_ have longer portions of their trajectories aligned with the chemotactic gradient. Remarkably, our experimental data suggest that 3-D migrating neutrophils maintain their chemotactic efficiency as ξ is decreased by varying their bias distance in concert with their persistence so that *s*_*θ*_ ≈ 3 *s*_*p*_. Thus, neutrophils seem to have developed a 3-D migration strategy that utilizes obstacles in the matrix as pivoting points to progressively reorient cell motion towards the chemoattractant source. In this strategy, denser matrices present more obstacles but also more pivoting points per unit volume, allowing cells to maintain efficient chemotaxis for a wide range of collagen densities. This behavior we found in 3-D contrasts with previous reports that neutrophils chemotaxing on 2-D surfaces only need one turn to correct their misalignment with respect to the gradient [53].

We tested our hypothesis about turning events and chemotactic efficiency by inhibiting Arp2/3, a complex shown to play a role in pathfinding during 3-D neutrophil migration [11]. Neutrophils migrating through straight microchannels have been reported to increase their speed and persistence after Arp2/3 inhibition [60]. However, the microchannels in those experiments did not challenge the cells to navigate physical barriers in search of migratory paths, which is different from our system and in many physiologically relevant contexts [57]. The neutrophils in our study had lower speeds after Arp2/3 inhibition due to their encounters with narrow matrix pores and the difficulties of either searching for alternate paths or exerting traction forces to expand the pores. Consistent with these results, previous work has shown that the Arp2/3 complex plays an important role in traction force generation for cancer cells [61]. Furthermore, our statistical analysis of trajectory orientation suggests a crucial role for the Arp2/3 complex in coordinating consecutive turns to circumvent narrow matrix pores, while keeping long-range directional motion towards the chemoattractant source. Specifically, Arp2/3 inhibited cells experienced a marked decrease in *s*_*p*_ /*s*_*θ*_, implying that they needed to concatenate a large number of turning events to correct misalignments in their motion. The effects of Arp2/3 inhibition were more prominent in high-[col] matrices, where neutrophils were more likely to encounter narrow, rigid pores. These results are consistent with previous studies involving dendritic cells and neutrophils, in which lamellipodia formation appeared to be crucial for navigational decision-making through complex geometries, especially as the microenvironment becomes more restrictive [17].

Recent studies with dendritic cells have shown that myosin-II becomes increasingly important during 3-D neutrophil migration as matrix pore sizes became smaller [16, 62]. In particular, the transition of the nucleus through matrix pores has been found to be a rate limiting factor in 3-D migration [62]. Even after myosin-II inhibition, neutrophils revealed their remarkable migratory versatility by increasing the rate of turning to circumvent matrix pores they would have otherwise been able to physically expand or squeeze through via contractile forces. While this adaptation translated into a decrease in cell speed, it did not seem to interfere with the ability of neutrophils to perform chemotaxis.

In conclusion, we analyzed the interplay between matrix deformation and turning events in 3-D neutrophil migration through matrices of varying pore sizes. We found that, as collagen concentration increases, neutrophils transition from a migratory regime characterized by transient large matrix deformations and straight trajectories to a second regime characterized by low matrix deformation and frequent turning events. More frequent turning translated into slower cell speed at higher collagen concentrations; however, it did not cause a decrease in chemotactic efficiency because the cells coordinated their more frequent turns to quickly resume directed motion. By pharmacologically inhibiting myosin-II contractility and Arp2/3 mediated pseudopod protrusion, we were able to trigger the transition to turning based migration at lower collagen concentrations than in untreated cells, confirming that this transition takes place when cells become unable to significantly deform the matrix. In addition, Arp2/3 inhibition caused a decrease in the frequency of turning events, leading to more persistent trajectories. However, it also caused a loss of coherence among turning events that made the Arp2/3-inhibited cells less efficient at chemotaxis. Overall, this work contributes to an enhanced mechanistic understanding of the role that the Arp 2/3 complex, myosin-II, and surrounding collagen density play in 3-D neutrophil chemotaxis, which in turn may provide crucial insights into molecules and processes to target for regulating pathological inflammatory responses [63].

## References

1. Aman, A. and T. Piotrowski, Cell migration during morphogenesis. Developmental biology, 2010. 341(1): p. 20–33.

2. Folkman, J., Angiogenesis, in Biology of endothelial cells. 1984, Springer. p. 412–428.

3. Hanahan, D. and R.A. Weinberg, Hallmarks of cancer: the next generation. cell, 2011. 144(5): p. 646–674.

4. Kolaczkowska, E. and P. Kubes, Neutrophil recruitment and function in health and inflammation. Nature Reviews Immunology, 2013. 13(3): p. 159.

5. Ley, K., et al., Getting to the site of inflammation: the leukocyte adhesion cascade updated. Nature Reviews Immunology, 2007. 7(9): p. 678.

6. Lam, P.-y. and A. Huttenlocher, Interstitial leukocyte migration in vivo. Current opinion in cell biology, 2013. 25(5): p. 650–658.

7. Wolf, K., et al. Collagen-based cell migration models in vitro and in vivo. in Seminars in cell & developmental biology. 2009. Elsevier.

8. Weninger, W., M. Biro, and R. Jain, Leukocyte migration in the interstitial space of non-lymphoid organs. Nature Reviews Immunology, 2014. 14(4): p. 232.

9. Sorokin, L., The impact of the extracellular matrix on inflammation. Nature Reviews Immunology, 2010. 10(10): p. 712.

10. Lämmermann, T. and M. Sixt, Mechanical modes of ‘amoeboid’cell migration. Current opinion in cell biology, 2009. 21(5): p. 636–644.

11. Fritz-Laylin, L.K., et al., Actin-based protrusions of migrating neutrophils are intrinsically lamellar and facilitate direction changes. Elife, 2017. 6: p. e26990.

12. Krause, M. and A. Gautreau, Steering cell migration: lamellipodium dynamics and the regulation of directional persistence. Nature reviews Molecular cell biology, 2014. 15(9): p. 577.

13. Weiner, O.D., et al., Spatial control of actin polymerization during neutrophil chemotaxis. Nature cell biology, 1999. 1(2): p. 75.

14. Krendel, M. and M.S. Mooseker, Myosins: tails (and heads) of functional diversity. Physiology, 2005. 20(4): p. 239–251.

15. Kaiser, C.A., et al., Molecular cell biology. 2007: WH Freeman.

16. Barbier, L., et al., Myosin II Activity Is Selectively Needed for Migration in Highly Confined Microenvironments in Mature Dendritic Cells. Frontiers in Immunology, 2019. 10: p. 747–747.

17. Leithner, A., et al., Diversified actin protrusions promote environmental exploration but are dispensable for locomotion of leukocytes. Nature cell biology, 2016. 18(11): p. 1253.

18. Renkawitz, J., et al., Nuclear positioning facilitates amoeboid migration along the path of least resistance. Nature, 2019. 568(7753): p. 546.

19. Manley, H.R., M.C. Keightley, and G.J. Lieschke, The neutrophil nucleus: an important influence on neutrophil migration and function. Frontiers in immunology, 2018. 9: p. 2867.

20. Salvermoser, M., et al., Nuclear deformation during neutrophil migration at sites of inflammation. Frontiers in Immunology, 2018. 9: p. 2680.

21. Wolf, K., et al., Physical limits of cell migration: control by ECM space and nuclear deformation and tuning by proteolysis and traction force. J Cell Biol, 2013. 201(7): p. 1069–1084.

22. Parkhurst, M.R. and W.M. Saltzman, Quantification of human neutrophil motility in three-dimensional collagen gels. Effect of collagen concentration. Biophysical journal, 1992. 61(2): p. 306–315.

23. Lammermann, T., et al., Rapid leukocyte migration by integrin-independent flowing and squeezing. Nature, 2008. 453(7191): p. 51–5.

24. Wu, P.-H., et al., Three-dimensional cell migration does not follow a random walk. Proceedings of the National Academy of Sciences, 2014. 111(11): p. 3949–3954.

25. Fraley, S.I., et al., Three-dimensional matrix fiber alignment modulates cell migration and MT1-MMP utility by spatially and temporally directing protrusions. Scientific reports, 2015. 5: p. 14580.

26. Millius, A. and O.D. Weiner, Chemotaxis in neutrophil-like HL-60 cells, in Chemotaxis. 2009, Springer. p. 167–177.

27. Afonso, P.V., et al., Discoidin domain receptor 2 regulates neutrophil chemotaxis in 3D collagen matrices. Blood, 2013. 121(9): p. 1644–1650.

28. Sixt, M. and T. Lämmermann, In vitro analysis of chemotactic leukocyte migration in 3D environments, in Cell Migration. 2011, Springer. p. 149–165.

29. Geraldo, S., A. Simon, and D.M. Vignjevic, Revealing the cytoskeletal organization of invasive cancer cells in 3D. Journal of visualized experiments: JoVE, 2013(80).

30. Schindelin, J., et al., Fiji: an open-source platform for biological-image analysis. Nature methods, 2012. 9(7): p. 676.

31. MATLAB, version 9.5 (R2018b). 2018, Natick, Massachusetts: The Mathworks Inc.

32. Molteni, M., et al., Fast two-dimensional bubble analysis of biopolymer filamentous networks pore size from confocal microscopy thin data stacks. Biophysical journal, 2013. 104(5): p. 1160–1169.

33. Münster, S. and B. Fabry, A simplified implementation of the bubble analysis of biopolymer network pores. Biophysical journal, 2013. 104(12): p. 2774–2775.

34. Raffel, M., et al., Particle image velocimetry: a practical guide. 2018: Springer.

35. del Alamo, J.C., et al., Three-dimensional quantification of cellular traction forces and mechanosensing of thin substrata by fourier traction force microscopy. PLoS One, 2013. 8(9): p. e69850.

36. Serrano, R., et al., Three-dimensional Monolayer Stress Microscopy. Biophysical journal, 2019.

37. Ayachit, U., The Paraview Guide: A Parallel Visualization Application. Kitware, 2015(978-1930934306).

38. Butler, J.P., et al., Traction fields, moments, and strain energy that cells exert on their surroundings. American Journal of Physiology-Cell Physiology, 2002. 282(3): p. C595–C605.

39. Fratzl, P., Collagen: structure and mechanics, an introduction, in Collagen. 2008, Springer. p. 1–13.

40. Marasco, W., et al., Purification and identification of formyl-methionyl-leucyl-phenylalanine as the major peptide neutrophil chemotactic factor produced by Escherichia coli. Journal of Biological Chemistry, 1984. 259(9): p. 5430–5439.

41. Hall, M.S., et al., Fibrous nonlinear elasticity enables positive mechanical feedback between cells and ECMs. Proceedings of the National Academy of Sciences, 2016. 113(49): p. 14043–14048.

42. Steinwachs, J., et al., Three-dimensional force microscopy of cells in biopolymer networks. Nature methods, 2015. 13(2): p. 171.

43. Nyberg, K.D., et al., Quantitative deformability cytometry: rapid, calibrated measurements of cell mechanical properties. Biophysical journal, 2017. 113(7): p. 1574–1584.

44. Kuntz, R.M. and W.M. Saltzman, Neutrophil motility in extracellular matrix gels: mesh size and adhesion affect speed of migration. Biophysical Journal, 1997. 72(3): p. 1472–1480.

45. Stroka, K.M. and H. Aranda-Espinoza, Neutrophils display biphasic relationship between migration and substrate stiffness. Cell Motil Cytoskeleton, 2009. 66(6): p. 328–41.

46. Petrie, R.J., A.D. Doyle, and K.M. Yamada, Random versus directionally persistent cell migration. Nature reviews Molecular cell biology, 2009. 10(8): p. 538.

47. Boal, D., Mechanics of the Cell. 2002.

48. Nolen, B., et al., Characterization of two classes of small molecule inhibitors of Arp2/3 complex. Nature, 2009. 460(7258): p. 1031.

49. Goley, E.D. and M.D. Welch, The ARP2/3 complex: an actin nucleator comes of age. Nature reviews Molecular cell biology, 2006. 7(10): p. 713.

50. Nourshargh, S. and R. Alon, Leukocyte migration into inflamed tissues. Immunity, 2014. 41(5): p. 694–707.

51. Kovács, M., et al., Mechanism of blebbistatin inhibition of myosin II. Journal of Biological Chemistry, 2004. 279(34): p. 35557–35563.

52. Straight, A.F., et al., Dissecting temporal and spatial control of cytokinesis with a myosin II Inhibitor. Science, 2003. 299(5613): p. 1743–1747.

53. Nossal, R. and S.H. Zigmond, Chemotropism indices for polymorphonuclear leukocytes. Biophysical Journal, 1976. 16(10): p. 1171–1182.

54. Schienbein, M. and H. Gruler, Langevin equation, Fokker-Planck equation and cell migration. Bulletin of Mathematical Biology, 1993. 55(3): p. 585–608.

55. Stout, D.A., et al., Mean deformation metrics for quantifying 3D cell–matrix interactions without requiring information about matrix materia. Proceedings of the National Academy of Sciences, 2016. 113(11): p. 2898–2903.

56. Handorf, A.M., et al., Tissue stiffness dictates development, homeostasis, and disease progression. Organogenesis, 2015. 11(1): p. 1–15.

57. Weigelin, B., G.-J. Bakker, and P. Friedl, Intravital third harmonic generation microscopy of collective melanoma cell invasion: principles of interface guidance and microvesicle dynamics. IntraVital, 2012. 1(1): p. 32–43.

58. Zaman, M.H., et al., Migration of tumor cells in 3D matrices is governed by matrix stiffness along with cell-matrix adhesion and proteolysis. Proceedings of the National Academy of Sciences, 2006. 103(29): p. 10889–10894.

59. Aung, A., et al., 3D traction stresses activate protease-dependent invasion of cancer cells. Biophysical journal, 2014. 107(11): p. 2528–2537.

60. Wilson, K., et al., Mechanisms of leading edge protrusion in interstitial migration. Nature communications, 2013. 4: p. 2896.

61. Giri, A., et al., The Arp2/3 complex mediates multigeneration dendritic protrusions for efficient 3-dimensional cancer cell migration. The FASEB Journal, 2013. 27(10): p. 4089–4099.

62. Davidson, P.M., et al., Nuclear deformability constitutes a rate-limiting step during cell migration in 3-D environments. Cellular and molecular bioengineering, 2014. 7(3): p. 293–306.

63. Mackay, C.R., Moving targets: cell migration inhibitors as new antiinflammatory therapies. Nature immunology, 2008. 9(9): p. 988.

