## Supplementary Information for "A balance between matrix deformation and the coordination of turning events governs directed neutrophil migration in 3-D matrices"

**Neutrophil turning model**

To further investigate whether the dynamics of turning of the chemotaxing neutrophils in our experiments, we constructed a biased random walk model [1]. This theoretical cell migration model describes cells with velocity magnitude $V$, moving in a chemotactic field with source at $y= \infty$(see Figure SI 1A). Consistent with the notation in the main text of the paper, the angle made by the velocity vector with the *y* axis (chemotactic direction) is denoted as $\theta$, and its evolution is given by the stochastic differential equation,

$d\theta\left( t \right)= -\gamma_{T}\sin\theta\left( t \right)dt+\Gamma\left( t \right)dt$,	[SI 1]

where $\gamma_{T}$ represents the response frequency (*i.e.*, inverse timescale) to correct the misalignment in a chemotactic field. The stochastic function $\Gamma(t)$ denotes the fluctuations in the turning response to model turning events to circumvent obstacles in the 3-D matrix, imperfections in the chemotactic signal transduction, and the inherent stochastic activity of the cell. We assume white noise for fluctuations, *i.e.,* $\Gamma\left( t \right)dt=\sqrt{\gamma_{p}} d\tilde{W}t$, where $\tilde{W}\left( t \right)$ is the standard Wiener process with Normal distribution$d\tilde{W}t=\sqrt{dt} N\left( 0,1 \right).$ The migratory persistence of the cells is quantified by $\gamma_{p}$, which is a frequency that characterizes the width of the chemotactic angle $(\theta)$distribution. The evolution of such distribution, $p\left( \theta,t \right)$ is governed by the associated Fokker-Planck equation:

$\frac{\partial p(\theta, t)}{\partial t}=\frac{\partial}{\partial\theta}\left[ \gamma_{T}\sin\theta\left( t \right) p\left( \theta,t \right) \right]+\frac{\gamma_{p}}{2} \frac{\partial^{2}}{\partial\theta^{2}} p\left( \theta,t \right).$	[SI2]

Scaling the timescale of the evolution equations with the persistence parameter, $\tau= t \gamma_{p} / 2$, we obtain the following non-dimensional system:

$d\theta\left( \tau\right)= -\lambda\sin\theta\left( \tau\right)d\tau+\sqrt{2 d\tau} N\left( 0,1 \right),$	[SI3]

$\frac{\partial p(\theta, \tau)}{\partial\tau}=\frac{\partial}{\partial\theta}\left[ \lambda\sin\theta\left( t \right) p\left( \theta,t \right) \right]+ \frac{\partial^{2}}{\partial\theta^{2}} p\left( \theta,t \right),$	[SI4]

where the non-dimensional parameter $\lambda= 2\frac{\gamma_{T}}{\gamma_{p}}$ is a key non-dimensional parameter that quantifies the ratio between the frequencies of biased and random turning. Following [1] for a similar system, the stationary solution to the Fokker-Planck equation is written as,

$p_{st}\left( \theta\right)=\frac{\mathcal{e}^{\lambda\cos\theta}}{\int_{0}^{2\pi} \mathcal{e}^{\lambda cos\theta}d\theta}$.	[SI5]

Using [1]’s definition of order parameter, we characterize the turning response curve of neutrophils with chemotactic order parameter defined as,

$<\cos\theta>=\int_{0}^{2\pi} p_{st}\left( \theta\right)\cos\theta d\theta= \frac{I_{1}\left( \lambda\right)}{I_{0}(\lambda)}$,	[SI6]

where $I_{0}$ and $I_{1}$ are modified Bessel’s functions. For the case of constant velocity magnitude $V$, temporal and spatial scales can be trivially interchanged given that $\Delta s=V\Delta t$, and the dimensionless parameter $\lambda$ can be expressed as $2\frac{\gamma_{T}}{\gamma_{p}}=2\frac{S_{p}}{S_{T}}$, where $S_{p}$ is the persistence length of the cells’ trajectories and $S_{T}$ is the lengthscale of biased turning, which is closely related to our experimentally determined $S_{\theta}.$

We plotted conditional probability distributions of p($\Delta\theta|\theta,\Delta s)$ for two different values of spatial separation along the trajectory, $\Delta s=S_{p}$and $\Delta s={5 S}_{p}$, similar to Figures 4E-F, Figure 5H, and Figure 6H. Figure SI1 B shows p($\Delta\theta\left| \theta,\Delta s \right)$ for $\lambda=0.3,$ which lies within the experimental range of $S_{p}/S_{\theta}$ found in our experiments for untreated and myosin-II treated chemotaxing cells (see Figure 7). The model results show good qualitative agreement with the experimental data from these cells (Figures 4E-F and 6H). A similar agreement is obtained when we plot p($\Delta\theta|\theta,\Delta s)$ for $\lambda=0.1$to model ck666-treated cells in restrictive matrices (Figure 5H, right panel). Thus, the directional dynamics of the neutrophils chemotaxing in 3-D matrices can be approximately represented by a biased random walk model for a wide range of matrix densities and pharmacological interventions.

**Supplementary Figure 1. Biased Random Walk Model. A)** Maps of probability distribution of $\Delta\theta$ conditional to the orientation$\theta$ with respect to the gradient $p\left( \Delta\theta|\theta,\Delta s \right)$**,** obtained from a biased random walk model with constant velocity and $\lambda= 2\frac{\gamma_{T}}{\gamma_{p}}=0.3$, which models cells with an efficient balance of persistence and chemotactic bias. The left and right panels display respectively $p\left( \Delta\theta|\theta,\Delta s=S_{p} \right)$ and $p\left( \Delta\theta|\theta,\Delta s={5 S}_{p} \right)$ vs. $\Delta\theta$ and $\theta$. **B)** Maps of $p\left( \Delta\theta|\theta,\Delta s \right)$**,** obtained from the same model with $\lambda= 2\frac{\gamma_{T}}{\gamma_{p}}=0.1$, which represents cells with an inefficient balance of persistence and chemotactic bias. The solid lines in panels **A-B** indicate $\theta={\Delta\theta}_{m}(\theta$) where ${\Delta\theta}_{m}$ is the mode of the $\Delta\theta$-distribution.

**Supplementary Table 1:** Number of cells used for cell trajectory based analyses in each experimental condition. These numbers reflect the number of cells after “non-motile” cells were filtered from larger data sets.

|  | **0.25 mg/mL** | **0.5 mg/mL** | **0.75 mg/mL** | **1 mg/mL** | **2mg/mL** |
| --- | --- | --- | --- | --- | --- |
| **no fMLP** | 202 | 643 | 644 | 576 | 17 |
| **fMLP, no treatment** | 1626 | 4238 | 3035 | 2873 | 1862 |
| **fMLP, ck666** | 426 | 1010 | 899 | 623 | 268 |
| **fMLP, blebbistatin** | 396 | 1105 | 240 | 323 | 243 |

**References**

1.	Schienbein, M. and H. Gruler, *Langevin equation, Fokker-Planck equation and cell migration.* Bulletin of Mathematical Biology, 1993. **55**(3): p. 585-608.
