## Supplementary figures and images for "A balance between matrix deformation and the coordination of turning events governs directed neutrophil migration in 3-D matrices"

### Supplementary Figure 1

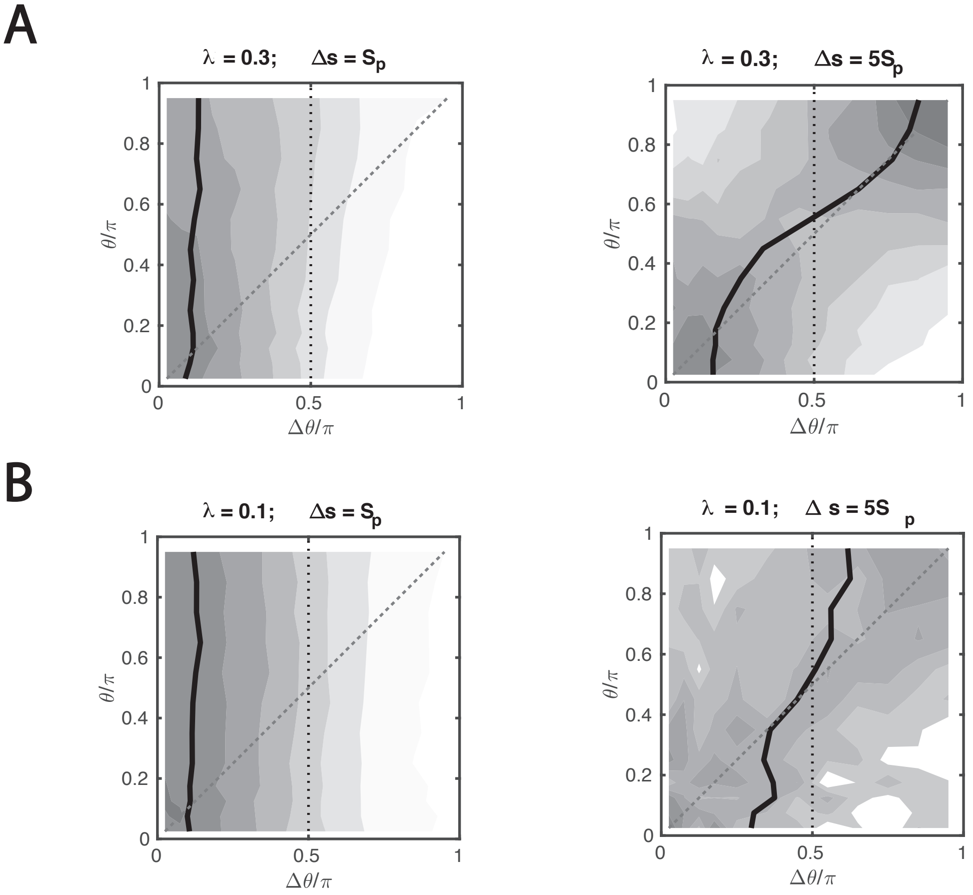
